# Low-level tuning biases in higher visual cortex reflect the semantic informativeness of visual features

**DOI:** 10.1101/2022.08.04.502850

**Authors:** Margaret Henderson, Michael J. Tarr, Leila Wehbe

## Abstract

Representations of visual and semantic information can overlap in human visual cortex, with the same neural populations exhibiting sensitivity to low-level features (orientation, spatial frequency, retinotopic position), and high-level semantic categories (faces, scenes). It has been hypothesized that this relationship between low-level visual and high-level category neural selectivity reflects natural scene statistics, such that neurons in a given category-selective region are tuned for low-level features or spatial positions that are diagnostic of the region’s preferred category. To address the generality of this “natural scene statistics” hypothesis, as well as how well it can account for responses to complex naturalistic images across visual cortex, we performed two complementary analyses. First, across a large set of rich natural scene images, we demonstrated reliable associations between low-level (Gabor) features and high-level semantic dimensions (indoor-outdoor, animacy, real-world size), with these relationships varying spatially across the visual field. Second, we used a large-scale fMRI dataset (the Natural Scenes Dataset) and a voxelwise forward encoding model to estimate the feature and spatial selectivity of neural populations throughout visual cortex. We found that voxels in category-selective visual regions exhibit systematic biases in their feature and spatial selectivity which are consistent with their hypothesized roles in category processing. We further showed that these low-level tuning biases are largely independent of viewed image category. Together, our results are consistent with a framework in which low-level feature selectivity contributes to the computation of high-level semantic category information in the brain.

## Introduction

Cortical responses to visual inputs demonstrate organization according to both high-level and low-level stimulus properties. High-level information about images, such as their membership in semantic categories, is reflected in the activation of spatially localized areas of the ventral visual cortex selective for categories such as faces, body parts, places, food, and words [1, 2, 3, 4, 5, 6, 7, 8]. At the same time, low- and mid-level visual features also elicit topographically regular patterns of activation in visual cortex, such as retinotopic maps of spatial position [9, 10, 11] and large-scale maps of selectivity for orientation [12, 13, 14], spatial frequency [15, 16], color [17, 18], and curvature [19, 20]. Given the hierarchical nature of processing in the visual system, understanding the relationship between selectivity for features at these different levels of complexity is critical for explicating the neural mechanisms by which high-level category information is computed in the brain.

Past work has suggested that low-level and high-level selectivity may be intertwined in their neural organization, in that category-selective visual regions exhibit systematic biases toward particular low-level properties of the visual environment. Not surprisingly, the low-level visual features most strongly represented in a given category-selective region tend to reflect the image statistics of the category in question. For example, scene-selective cortical regions, such as the parahippocampal place area (PPA), have been shown to be more responsive to cardinal (vertical and horizontal) orientations and rectilinear contour features than diagonal orientations and curved contours [21, 22, 23]. PPA has also been shown to be biased toward high spatial frequencies over low [24]. In contrast, areas of the visual system selectively responsive to faces have been shown to overlap with selectivity for curved features [25, 19, 20], and these face-selective areas may be more responsive to low spatial frequencies [24]. In addition to feature selectivity, spatial selectivity also appears to co-vary with category responsiveness, with face-selective and word-selective cortical regions tending to have biases toward the central visual field, and scene-selective cortical regions tending to have more peripheral biases [26, 27].

A unifying explanation for these findings is based on the observation that, as mentioned, for many categories of natural images, the statistics of low-level visual features differ depending on the semantic content of images [28, 29]. For example, large-scale outdoor scene images are dominated by horizontal orientations and high spatial frequencies, while close-up images of objects may have a more isotropic distribution of orientations and a more energy at low spatial frequencies [28]. Beyond spectral features, mid-level features like the overall curvature in an image may co-vary with object-level distinctions such as real-world size and animacy [30, 31]. Such statistical associations may be reflected in the organization of visual cortex due to learning – either over the course of evolution or during an individual’s lifetime. These observations suggest that category-selective visual regions follow a principle whereby they are biased in favor of low-level image properties that are informative for their “preferred” category, and these biases may play a functional role in both learning and categorization [32, 33].

Supporting the hypothesis that image statistics constrain the low-level biases found in category-selective visual cortex, the selectivity for cardinal orientations and rectilinear contours in scene-selective visual areas has been related to the fact that scene stimuli contain more cardinal orientations and rectilinear angles than non-scene stimuli [22]. A similar idea may hold for color selectivity, based on the finding that color tuning of neurons in ventral object-selective cortex is biased in favor of colors that are associated with objects [34]. Biases in spatial coverage of the visual field may also be understood in this framework, for example, the foveal eccentricity biases found in face- and word-selective cortical regions may be related to the use of high spatial frequency information in identifying these classes of stimuli, and the fact that they tend to be foveated, leading to an association with the central visual field [27, 26].

While these results provide some insight into the origins of low-level biases in visual cortex, much of the supporting work has focused on a small range of visual stimulus classes, often using controlled synthetic stimuli or objects on isolated backgrounds rather than natural scenes [22, 26, 27]. In addition, there has been a tendency to focus on only one brain region or a small group of regions at a time [21, 22, 26, 27, 23]. As a result, the generality of this hypothesis is somewhat uncertain, particularly with respect to how well it can account for findings across a range of visual areas during naturalistic image viewing. As an alternative, it has been suggested that semantic category selectivity reflects representations at a high level of abstraction only [35]. That is, the critical dimension in the organization of high-level visual cortex is category membership in and of itself rather than selectivity for continuously varying low-level features that may be associated with each given category. On this view, low-level feature biases do not play a functional role in computing category membership, and, instead, may reflect more general structural constraints, such as inheritance of selectivity from an earlier area in the processing stream. The implications of this alternative are that low-level feature biases measured in category-selective visual areas may *not* correlate with the low-level features and/or positions that are most diagnostic of each area’s preferred category.

To distinguish between these alternatives, we provide two complementary sets of analyses. First, we take advantage of a large, richly-annotated natural scene image database [36] to demonstrate consistent associations between low-level features (orientation, spatial frequency) and high-level semantic category labels. Next, we use a recently released, large-scale open fMRI dataset collected by Allen et. al. [37] (the “Natural Scenes Dataset” or NSD) to show that these same associations are reflected in the patterns of feature and spatial selectivity estimated from neural population responses measured while human participants view natural scene images. Importantly, our use of naturalistic images allows us to demonstrate a link between neural response properties and natural scene statistics within an ecologically relevant setting, in contrast to past work with simpler stimuli. Our results also provide a proof-of-concept that low-level feature and spatial biases can be reliably measured in the human brain using complex, naturalistic stimuli. Furthermore, our use of a high-resolution, whole-brain fMRI dataset provides an opportunity to assess our hypothesis across the brain rather than in a limited region of cortex. Based on the convergence among our analyses, we argue that low-level feature and spatial biases may play a key role in the processing of semantic categories within high-level visual areas.

## Results

Our overall experimental goal was to measure the selectivity of cortical populations for visual features and spatial positions, and determine if the properties of selectivity in each visual area can be understood in terms of natural image statistics. We used a two-pronged approach to critically assess different hypotheses. First, we analyzed the relationship between low-level visual (Gabor) features and high-level semantic categories across a large set of natural scene images (Figure 1A). Second, we performed a comprehensive analysis of the orientation, spatial frequency, and spatial selectivity of neural populations throughout visual cortex. The measured neural activity in these populations was taken from the Natural Scenes Dataset (“NSD” [37]) in which fMRI scanning was performed while human observers viewed COCO images (an independent set from those used in our first analysis). We constructed voxel-wise predictive encoding models based on image-computable features and used these models to estimate voxel selectivity for low-level features as well as spatial position (Figure 1B). To examine the relationship between feature selectivity and category selectivity, we compared our results across early visual areas, dorsal visual areas, and a range of category-selective cortical regions and determined whether the specific biases detected in each area corresponded with that area’s presumed role in category processing.

**Figure 1:**
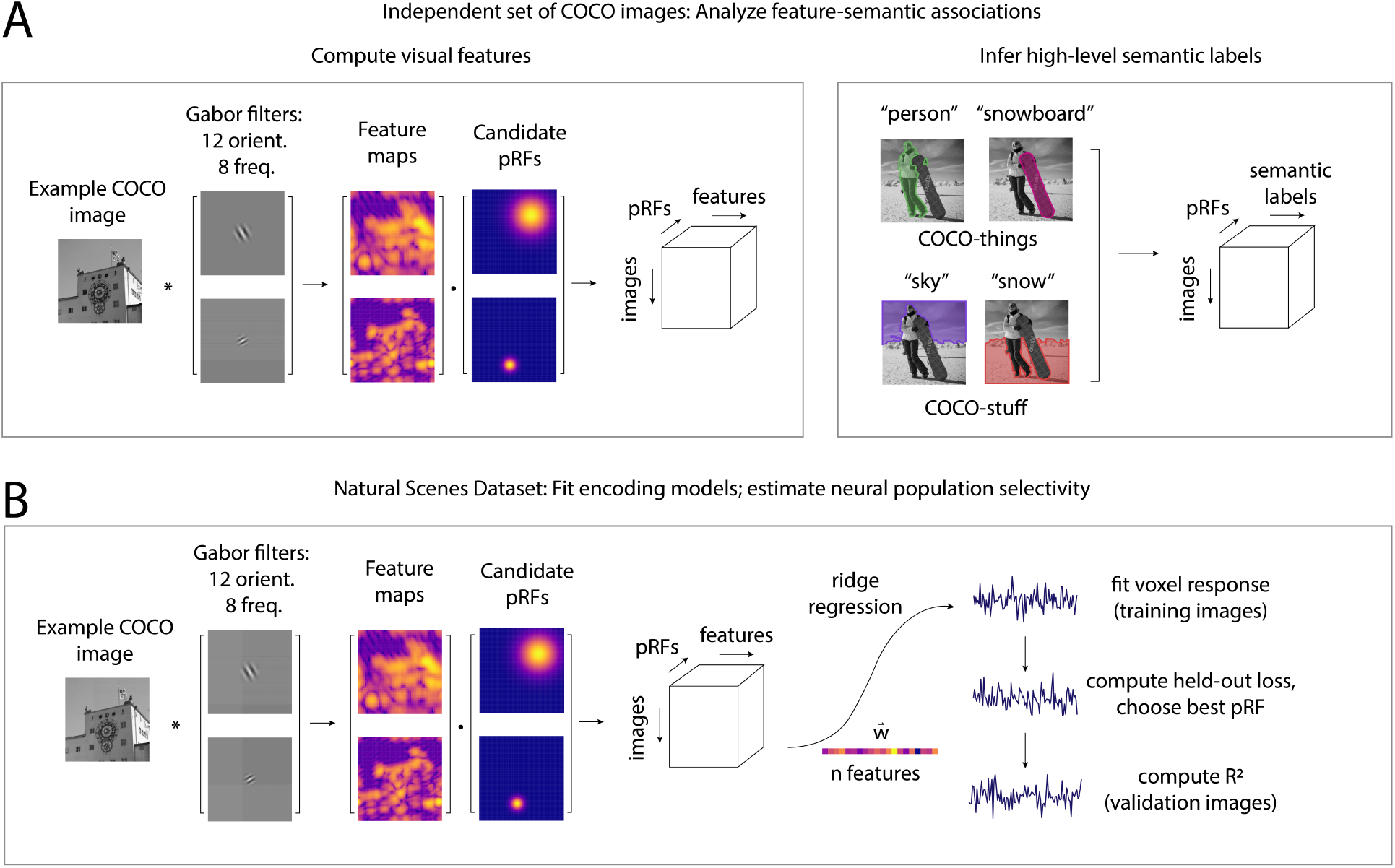
Overview of our two analysis procedures. **(A)** In the first set of analyses (results are presented in Figure 2) we analyzed the associations between Gabor features and high-level semantic labels across a set of 10,000 COCO images (note that no fMRI data is involved in this step). The left box depicts a schematic of the feature extraction procedure: for each natural scene image, we used a Gabor feature bank to compute a stack of feature maps, then applied various spatial weighting matrices (i.e., candidate population receptive fields/pRFs [38]) to compute the estimated activation in each feature map for a range of spatial positions and pooling field sizes (see *Methods: Gabor Features* for details). The right box depicts examples of the semantic category labels included in the COCO dataset[36]; each label is accompanied by a spatial mask indicating the corresponding object’s position in the image. We used these labels to infer high-level semantic category labels for each image, labeling either the entire image (indoor-outdoor) or labeling each pRF individually (animacy, realworld size); see *Methods: Semantic Features* for details. **(B)** In the second set of analyses (results are presented in Figure 3 - Figure 6), we used the Natural Scenes Dataset [37] to learn an encoding model predicting each fMRI voxel’s response as a weighted sum of image features in the voxel’s pRF. The feature extraction procedure was identical to **(A)**, but performed on a different set of images (the images viewed by each fMRI participant). To fit the model for each voxel, we first learned a set of weights for each candidate pRF (using ridge regression), then chose the pRF which resulted in the lowest loss on held-out data. See *Methods: Model Fitting Procedure* for more details. Note that while this figure illustrates use of a Gabor feature bank, we used the same approach for other feature spaces; see text for details.

### Associations Between Low-Level Features and High-Level Semantic Categories

Across a set of 10,000 natural scene images sampled from COCO, we observed several key relationships between low-level Gabor features and high-level semantic categories (Figure 2). Here we focus on three high-level semantic categories: indoor versus outdoor (a scene-level property), animacy, and real-world size (both object-level properties); see *Methods: Semantic Features* for details on how high-level category labels were determined. In the orientation domain, we found that images labeled as outdoor scenes, as well as image patches containing large, inanimate objects, were positively associated with horizontal (90°) orientations (Figure 2A, 2B). In contrast, image patches containing animate objects and/or objects with a small real-world size were positively associated with diagonal orientations (45°/135°). Correlations of each category with vertically oriented (0°) features varied across spatial frequency: at low spatial frequencies, vertical orientations were positively associated with indoor scenes and small objects, while at high spatial frequencies, vertical orientations were positively associated with outdoor scenes and large objects. When averaging over orientation and focusing on the spatial frequency axis itself, we found that higher spatial frequency features were more positively associated with outdoor scene images, and were also weakly associated with animate objects.

**Figure 2:**
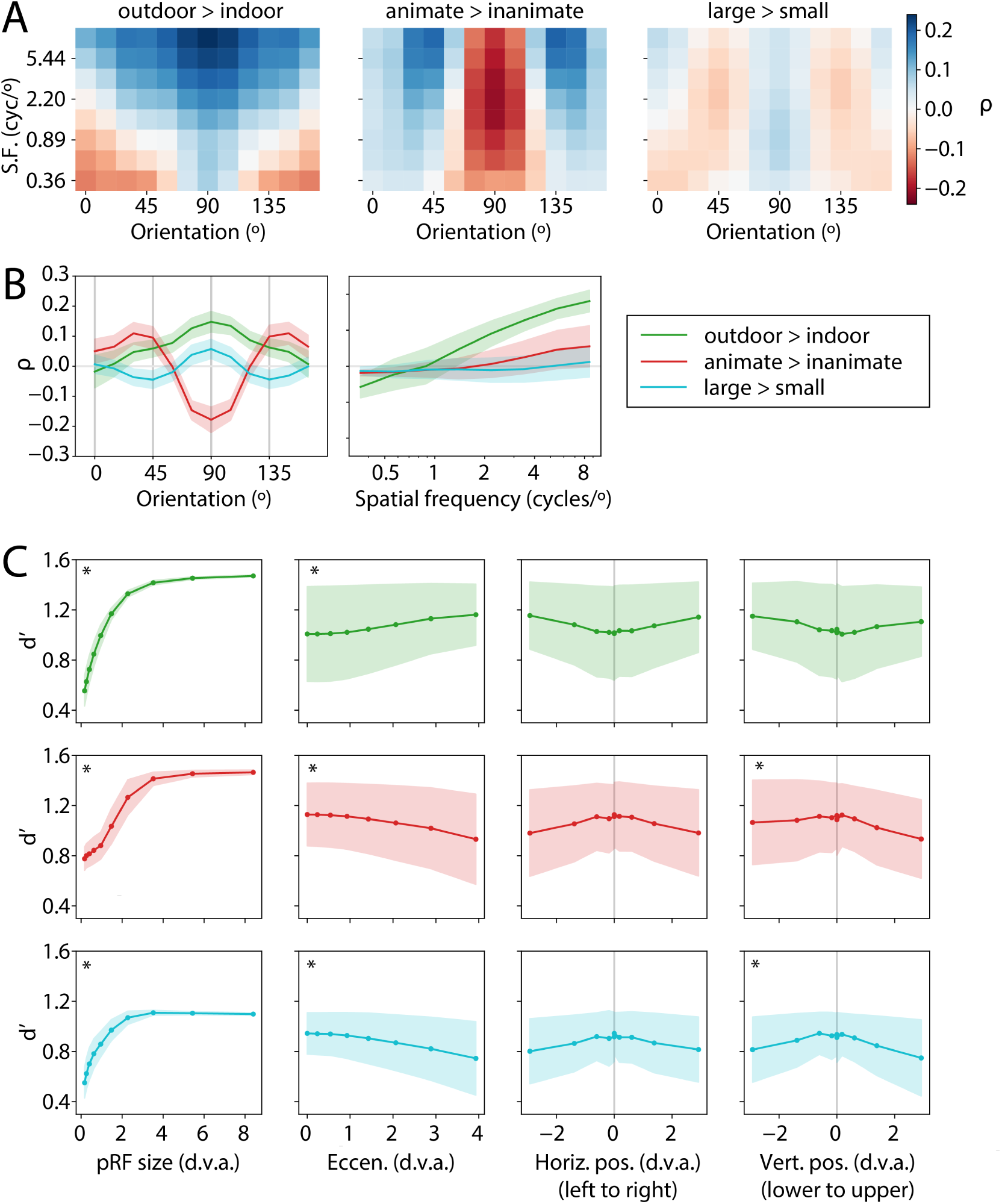
High-level semantic categories are each associated with distinct patterns of low-level visual features. **(A)** The partial correlation coefficient (*ρ*) for each Gabor feature channel with each semantic axis, with channels organized by spatial frequency on the y-axis and orientation on the x-axis. Our orientation axis is defined such that 0°=vertical and 90°=horizontal. **(B)** The partial correlation coefficients averaged over either spatial frequency (left panel) or orientation (right panel), with error bars indicating ± 1 s.d. across pRFs. **(C)** Cross-validated performance (d’) of a linear decoder at discriminating each category distinction based on patterns of activation across all 96 Gabor feature channels (see *Methods: Measuring Feature-Semantic Associations*). Rows represent different semantic axes (colors as in **(B)**), each column represents values binned according to one pRF parameter (d.v.a = degrees of visual angle). Error bars indicate ± 1 s.d. across pRFs within each parameter bin, * indicates a significant linear relationship evaluated using a permutation test, FDR corrected *α*=0.01. All analyses in this figure were performed using a set of 10,000 COCO images independent from the ones used in the NSD dataset; see *Methods: Measuring Feature-Semantic Associations* for details.

The observed statistical associations between Gabor features and high-level semantic categories indicate that low-level visual features may serve as informative cues for the detection of those categories. To further investigate this effect, we used a linear decoding analysis to examine how the information contained across all Gabor feature channels varied as a function of spatial position in the visual field. Since both our Gabor features and semantic labels were computed in a spatially-specific manner across a grid of candidate model pRFs, we performed decoding for each pRF separately and then examined how decoding performance (*d*^*′*^, see *Methods: Measuring Feature-Semantic Associations* for details) varied as a function of pRF size and position (Figure 2C). First, unsurprisingly, larger pRFs contained more information for all three semantic axes, with *d*^*′*^ saturating for each axis at a size of about 4 degrees of visual angle. Second, the effect of eccentricity differed across pRFs, with more peripheral pRFs containing more information about the indoor/outdoor axis, but more central pRFs containing more information about the animacy and real-world size distinctions. Third, comparing the upper and lower visual field, we found that pRFs in the lower visual field were more informative for determining animacy and real-world size than those in the upper visual field, but no effect of vertical position was found for the indoor-outdoor axis. No differences were detected between the left and right visual fields.

### Feature Selectivity of Neural Populations in Visual Cortex

Given this evidence for associations between image features and semantic categories, we next evaluated whether neural populations in human cortex exhibit biased tuning that reflects these associations. Specifically, we focused on voxels within several regions of interest (ROIs) in visual cortex: early retinotopic ROIs (V1, V2, V3, and hV4), dorsal visual ROIs (V3ab, IPS), place-selective ROIs (OPA, PPA, RSC), face-selective ROIs (OFA, FFA), and a body-selective ROI (EBA). All ROIs were defined using independent functional mapping data; see *Methods: Human Participants and Acquisition of fMRI Data* for details. The rationale for our choice of these regions is that the category-selective ROIs will allow us to test pre-existing hypotheses regarding the functional roles of these regions in category processing, while including early visual and dorsal ROIs will allow us to explore the generalizability of our findings beyond the typically explored category-selective regions of ventral visual cortex.

To measure selectivity for low-level visual features and spatial positions, we constructed a forward encoding model for each voxel that modeled its response to each image as a linear combination of a set of image-computable features. Our framework also incorporated a model of each voxel’s spatial selectivity or pRF (see Figure 1B and *Methods: Encoding Model Design* for details). We used two different visual feature spaces to construct encoding models. The first feature space was a set of Gabor features with channels corresponding to different combinations of orientation and spatial frequency; this model allowed us to assess voxel selectivity for easily interpretable low-level visual features. The Gabor encoding model was able to predict voxel responses to held-out images with good accuracy (R^2^) across a range of visual ROIs, with highest average R^2^ in V1, and performance generally declining in more anterior ROIs (Figure 3A). The second feature space was a set of features from a deep neural network (AlexNet; [39]); this model was used primarily to fit the pRF parameters for each voxel, and was chosen because it provided higher predictive performance than the Gabor model, particularly in higher levels of visual cortex (Figure 3A, see *Methods: Model Fitting Procedure* for more details).

**Figure 3:**
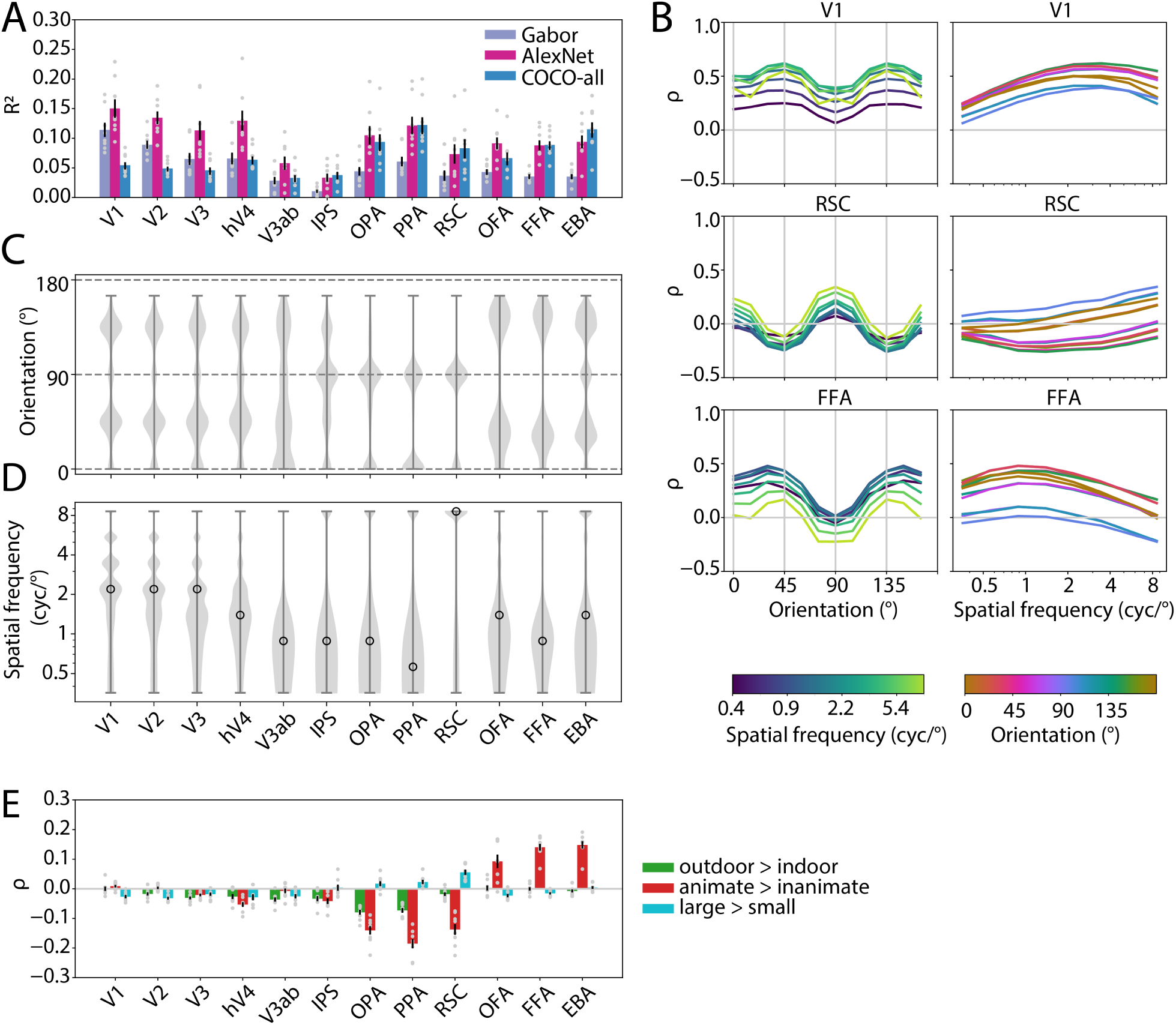
Feature selectivity of voxels differs across early visual and category-selective ROIs. **(A)** The average cross-validated accuracy (R^2^) of our encoding model. Colors correspond to different visual or semantic feature spaces, see *Methods: Feature Spaces* for details. The Gabor features (light purple bars) were used to estimate parameters in **(B-D)**, see text for discussion of AlexNet and COCO-all models. Each bar represents R^2^ averaged over voxels within one ROI; gray points indicate single participants, error bars represent ± 1 SEM across 8 participants. **(B)** The average orientation (left) and spatial frequency (right) sensitivity profile of voxels in several ROIs (averaged over 8 participants), with colored lines indicating different spatial frequency (or orientation) levels. Feature sensitivity (*ρ*) was estimated by computing the correlation between the predicted encoding model response for validation set images and the activation in each feature channel for the same images (see *Methods: Estimating Voxel Visual Feature Selectivity* for details). Supplementary Figure 1 and Supplementary Figure 2 show these profiles across all ROIs. **(C)** The distribution of “preferred” orientation for voxels in each ROI, where preferred orientation is defined as the orientation associated with the maximum sensitivity (across all frequencies). Dotted lines indicate the cardinal orientations (0°=vertical, 90°=horizontal). **(D)** The distribution of preferred spatial frequency for voxels in each ROI, where preferred frequency is defined as the frequency associated with the maximum sensitivity (across all orientations). Open circle indicates the median of the distribution. Values in **(C)** and **(D)** reflect voxels pooled across all 8 participants. **(E)** The partial correlation (*ρ*) between voxel responses in each ROI and three high-level semantic dimensions (see *Methods: Estimating Voxel Semantic Selectivity* for details), averaged over all voxels within each ROI and participant. Gray points indicate single participants, error bars represent ± 1 SEM across 8 participants.

Once each voxelwise encoding model was fit, we used the models to estimate voxel sensitivity for individual model features. Our measure of feature sensitivity was computed by generating encoding model predicted responses to images in the validation image set (i.e., a set of images not used during model training), and then correlating the predicted responses with the continuous activation values in each feature channel (see *Methods: Estimating Voxel Visual Feature Selectivity* for details). Since the orientation and spatial frequency of each Gabor feature channel is known, we then plotted the sensitivity values as a function of orientation to yield orientation sensitivity profiles (Figure 3B, left column, and Supplementary Figure 1) or as a function of spatial frequency to yield spatial frequency sensitivity profiles (Figure 3B, right column and Supplementary Figure 2). Plotting the feature sensitivity profiles, averaged across all voxels in each ROI, revealed several key differences among ROIs. First, early visual ROIs, while having positive sensitivity on average for all Gabor feature channels, had the highest sensitivity for oblique (45°/135°) orientations. This effect, which diverges from the common finding of a cardinal orientation bias in early visual cortex, may be related to the broad spatial frequency content of our natural image stimuli; we return to this issue in the Discussion. In contrast, place-selective regions tended to have largest average sensitivity for vertical (0°) and horizontal (90°) orientations. Face- and body-selective regions displayed a different pattern, with the highest sensitivity values for oblique orientations 30° and 150°. Along the spatial frequency axis, early visual areas tended to have the highest average sensitivity for spatial frequencies between 2-5 cycles/°, while spatial frequency sensitivity profiles in face-selective regions peaked at a lower spatial frequency of around 1-2 cycles/°. Place-selective region RSC showed maximal sensitivity for high spatial frequencies.

Consistent with the ROI-averaged sensitivity profiles, differences were also evident in the distribution of peak sensitivity values (i.e., “preferred” feature values) across individual voxels within each ROI (Figure 3C, 3D). With regard to orientation sensitivity, early visual, as well as face-selective and body-selective ROIs, tended to have mostly voxels preferring oblique orientations (45°/135°), although as in the previous analysis, the distribution in face-selective ROIs was shifted slightly toward vertical relative to the distribution in early visual cortex. In contrast, both IPS and the place-selective ROIs tended to have more voxels preferring the cardinal orientations (0°/90°). Intermediate dorsal area V3ab had a more uniform distribution of peak orientation sensitivity values that included a combination of oblique and cardinal orientations. In the spatial frequency dimension, early visual ROIs tended to have mostly voxels preferring mid-to-high spatial frequencies, with the median preferred spatial frequency tending to be lower in more anterior category-selective regions. The primary exception to this was RSC, where most of the voxels had their peak sensitivity at a higher spatial frequency. Several other ROIs, particularly EBA, OPA, PPA, and IPS, also had a secondary, smaller group of voxels preferring high spatial frequencies (i.e, top of violins in Figure 3D).

To further aid interpretation of these results and link them to our image statistics analysis (Figure 2), we also analyzed the degree to which voxels in each ROI were selective for each high-level semantic category distinction (Figure 3E). Unsurprisingly, this revealed a distinction between the face- and body-selective areas versus the place-selective areas, in that OFA, FFA, and EBA were consistently more activated for animate than inanimate objects, while RSC, PPA, and OPA were consistently more activated for inanimate objects. RSC additionally showed a consistent preference for large over small objects, which was also evident more weakly in OPA and PPA, while OFA and FFA each exhibited a slight preference for small over large objects. Taking these observations into account along with the feature sensitivity results, a consistent relationship is evident. On the one hand, place-selective ROIs are biased towards orientations (0°/90°) that match their selectivity for large, inanimate objects. On the other hand, face-selective and body-selective areas are biased toward oblique orientations, specifically 30°/150°, which are positively associated with animacy. Thus, these results support our prediction that category-selective visual areas exhibit biases toward low-level features diagnostic of their preferred category.

Of note, these results were not dependent on assuming any particular shape for orientation and frequency sensitivity profiles. A supplementary analysis (Supplementary Figure 3) revealed that while the spatial frequency sensitivity profiles for most voxels tended to have only one peak, a significant proportion of voxels across all areas had orientation sensitivity profiles that were bimodal, having two peaks (see *Methods: Counting Peaks in Feature Sensitivity Profiles* for details). When these bimodal voxels were analyzed based on the orientations at which their two peaks occurred, they largely fell into groups that matched the results of our previous analysis (i.e., bimodal voxels in face-selective areas tended to have two peaks at 30°/150°, bimodal voxels in place-selective areas tended to have two peaks at 0°/90°). This suggests that the orientation biases in these areas were widespread across voxels, regardless of the exact shape of their sensitivity profiles.

### Spatial Selectivity of Neural Populations in Visual Cortex

Analyzing the spatial selectivity of voxels estimated by our encoding model (pRFs; see *Methods: Encoding Model Design* for details) revealed several trends among brain regions. First, we found that the median size (*σ*) of pRFs was smallest in V1, and tended to increase progressively along the anterior axis of the brain, with the largest median pRFs sizes observed in PPA, RSC, and FFA (Figure 4A). Within each ROI, pRF sizes tended to scale approximately linearly with pRF eccentricity (Figure 4B). These results are consistent with past work [38, 40, 41], and thus provide validation of our model fitting procedure. Also consistent with past work, we found that in early visual areas, when our estimates of pRF angular position for each voxel were compared to our estimates of preferred orientation from the previous analysis (Figure 3C), a clear and consistent correlation was evident between preferred orientation and preferred angular position (Supplementary Figure 4). This result aligns with previous findings of radial bias in early visual cortex [13, 14], and provides additional support for the validity of both our spatial selectivity and feature selectivity estimates.

**Figure 4:**
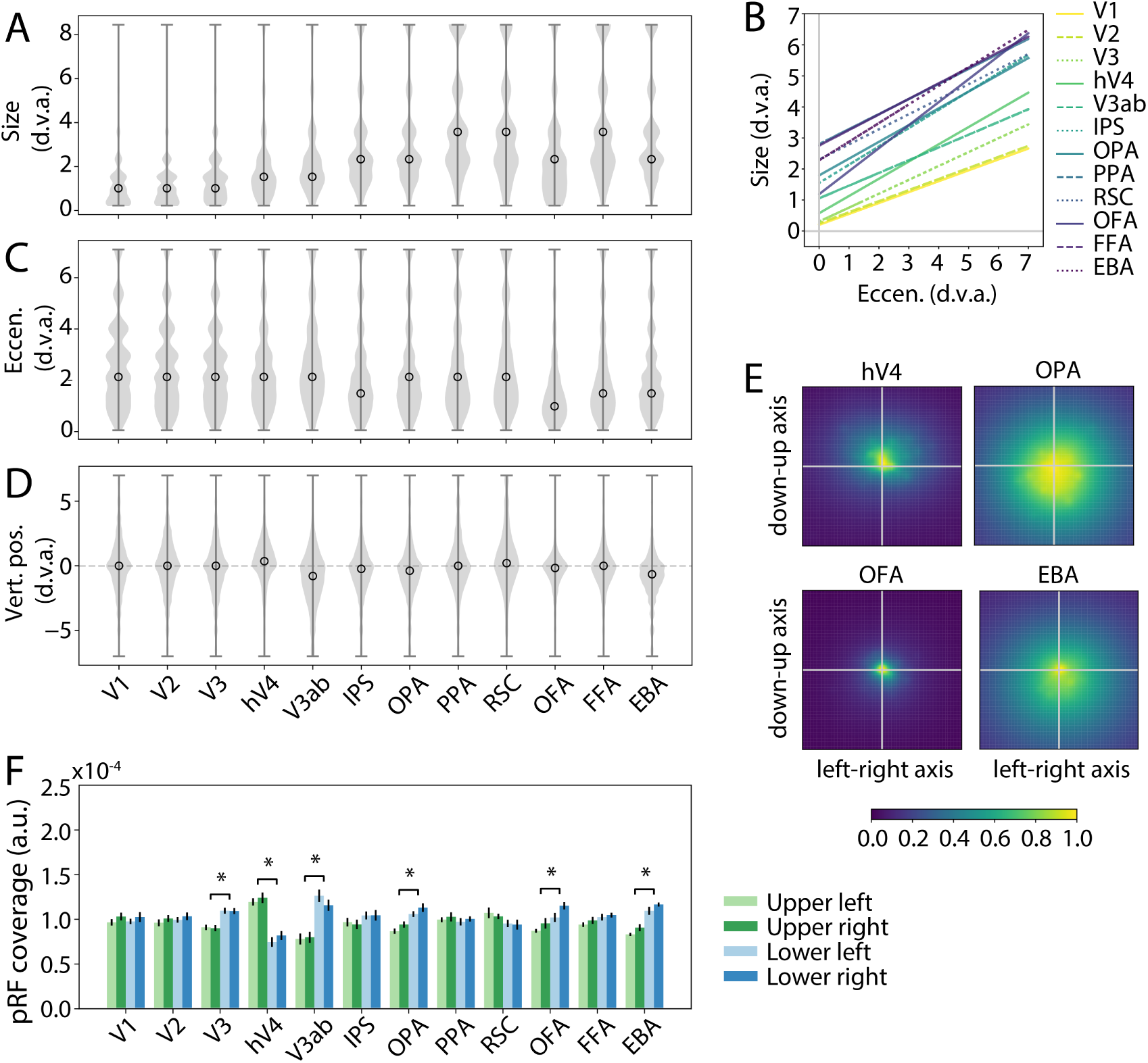
Spatial selectivity of voxels differs across early visual and category-selective ROIs. **(A)** The distribution of the best size (*σ*) parameter for all voxels in each ROI, combined across 8 participants. Open circle indicates the median. **(B)** The distribution of pRF eccentricity. **(C)** The distribution of pRF vertical position. See Supplementary Figure 5 for distributions of horizontal position and angular position. **(D)** The best linear fit lines for the relationship between size and eccentricity of pRFs across all voxels within each ROI, combining across all participants. **(E)** Visualization of the coverage of the visual field by pRFs in three example ROIs, obtained by averaging the pRFs over individual voxels in each ROI. **(F)** The mean of the aggregated pRF values in each visual field quadrant (i.e., mean of quadrants in each image of panel **(E)**). * indicates significance of paired *t*-test for the upper versus lower visual field difference, FDR corrected *α*=0.01. Error bars indicate ± 1 SEM across participants. Note that values in panel **(E)** have been normalized to have a maximum of 1 for visualization purposes, values in **(F)** are not normalized.

In addition to these general trends, further examination of the pRF parameters in each ROI indicated biases toward particular portions of the visual field. Focusing first on the eccentricity of pRF centers, we found that the median eccentricity was lowest in OFA, followed by FFA, EBA, and IPS, while place-selective regions had relatively more voxels with pRFs in the periphery (Figure 4C). This result is consistent with previous findings of a foveal bias for face representations and a peripheral bias for scene representations [26, 27]. In addition to eccentricity, different ROIs also exhibited differences in the distribution of pRF vertical positions (Figure 4D). Both hV4 and RSC tended to have more pRFs with centers in the upper visual field, while V3ab, OPA, and EBA each tended to have more centers concentrated in the lower visual field. To quantify this effect, we computed a measure of pRF coverage across the entire visual field, by aggregating all pRFs across all voxels for each participant and then taking the mean (Figure 4E; see *Methods: pRF Coverage Analysis*). Taking the average of these coverage plots within each visual field quadrant provided a measure of pRF coverage for each quadrant (Figure 4F), [42]. Note that this measure is different from counting the number of pRF centers in each quadrant because it takes into account pRF size as well as center. Across participants, a significant interaction between pRF coverage of the upper vs. lower visual field and ROI was found (three way repeated measures ANOVA with ROI, vertical position, and horizontal position as factors: main effect of ROI, *F*_(11,77)_=-2.33, *p*=1.0000; main effect of vertical position, *F*_(1,7)_=3.29, *p*=0.1124; main effect of horizontal position, *F*_(1,7)_=3.89, *p*=0.0893; interaction between ROI and vertical position, *F*_(11,77)_=10.68, *p <* 10^*−*4^; see Supplementary Table 5 for all interactions). Post-hoc tests revealed that pRF coverage was higher for the upper versus the lower visual field in hV4, while average pRF coverage was higher for the lower versus the upper visual field in V3, V3ab, OPA, OFA, and EBA (paired *t* -test with permutation; FDR corrected *α*=0.01). No differences in coverage of the left versus the right visual field were observed (see also Supplementary Figure 5A).

Comparing these spatial biases to the distribution of high-level semantic category information across the visual field (Figure 2C), a relationship is apparent. The half of the visual field containing the most information about the object properties animacy and real-world size (i.e., the lower visual field) is over-represented within several category-selective ROIs, OPA, OFA, and EBA. The strongest bias towards the central visual field, which, as with the lower visual field, was found to contain more information about animacy and real-world size, was found in face-selective area OFA. Thus, our analyses of spatial selectivity are consistent with the hypothesis that category-selective ROIs may be biased towards semantically-informative portions of the visual field.

### Separability of Feature Selectivity and Category Selectivity

Given that our image set consists of natural scene images conveying semantic category information in addition to visual features, an important question is the extent to which our measurements of visual feature selectivity are influenced by responses to the semantic information itself. For instance, if a voxel shows a preferential response to Category A, and Category A is correlated with feature B, it is possible that our feature selectivity analyses will show selectivity for feature B even if the voxel does not have any true feature selectivity and is better described as category-selective. To test for this possibility, we performed two additional analyses. First, we regressed out the contributions of explicit category selectivity from each voxel response by building an encoding model whose features described the presence of various categories in the image (see *Methods: Semantic Features* and *Methods: Regressing Out Category Selectivity* for details). We then re-fit the Gabor encoding model based on just the residuals of this model. The rationale for this is that any feature selectivity detected with this method likely reflects true feature selectivity, exceeding what is explainable based on category selectivity. Second, for each of our high-level semantic dimensions, we re-fit the encoding models using images of only one category label at a time (i.e., for the indoor-outdoor dimension, we separately analyzed the indoor scene images and the outdoor scene images; see *Methods: Fitting Models for Single Categories* for details). This analysis provided a complementary method for removing the effects of category selectivity from voxel responses. It also allowed us to test whether our results were robust to differences in the overall feature statistics of the image set – since images of different categories are likely to have different distributions of features (i.e., Figure 2), fitting the encoding model for different categories separately helped to determine whether these statistical differences influenced our measurements of feature selectivity.

Critically, feature selectivity remained largely consistent after removing the contributions of explicit semantic category selectivity from the voxel responses (Figure 5). Our category-based model alone did predict a substantial portion of voxel response variance (Figure 3A,“COCO-all”/blue bars), and as a result, re-fitting the Gabor model to the residuals of this categorical model resulted in a drop in R^2^ compared to fitting on the raw data, especially in category-selective regions (Figure 5A). Nevertheless, many voxels were still well-predicted by the Gabor model fit to the residuals (see Supplementary Table 4 for list of voxel counts after thresholding). Moreover, the feature sensitivity of these voxels was relatively consistent when compared to the feature sensitivity estimated from the raw data, as measured by a correlation coefficient between the feature sensitivity profiles across the two methods (Figure 5B). The largest changes in feature sensitivity when regressing out category selectivity were observed in PPA, followed by OPA, RSC, and EBA, although the correlation between feature sensitivity from the raw data and from the residuals was still positive on average in all areas. To understand these changes in feature selectivity, we visualized the distribution of peak orientation and peak spatial frequency across voxels in all category-selective ROIs (Figure 5C), and plotted orientation sensitivity profiles for OPA, PPA, and RSC (Figure 5D). These data suggest that changes in feature selectivity in place-selective areas were primarily related to a change in the spatial frequency sensitivity of voxels. This shift was most dramatic in PPA, where the distribution of peak spatial frequency across voxels shifted from being more concentrated at low spatial frequencies for the raw data, to more concentrated at high spatial frequencies for the residual data. In contrast, in face-selective areas OFA and FFA these analyses established that the distributions of peak orientations and peak spatial frequencies were consistent whether estimated from the raw data or the category model residuals, suggesting that feature selectivity in these areas was largely independent of category selectivity.

**Figure 5:**
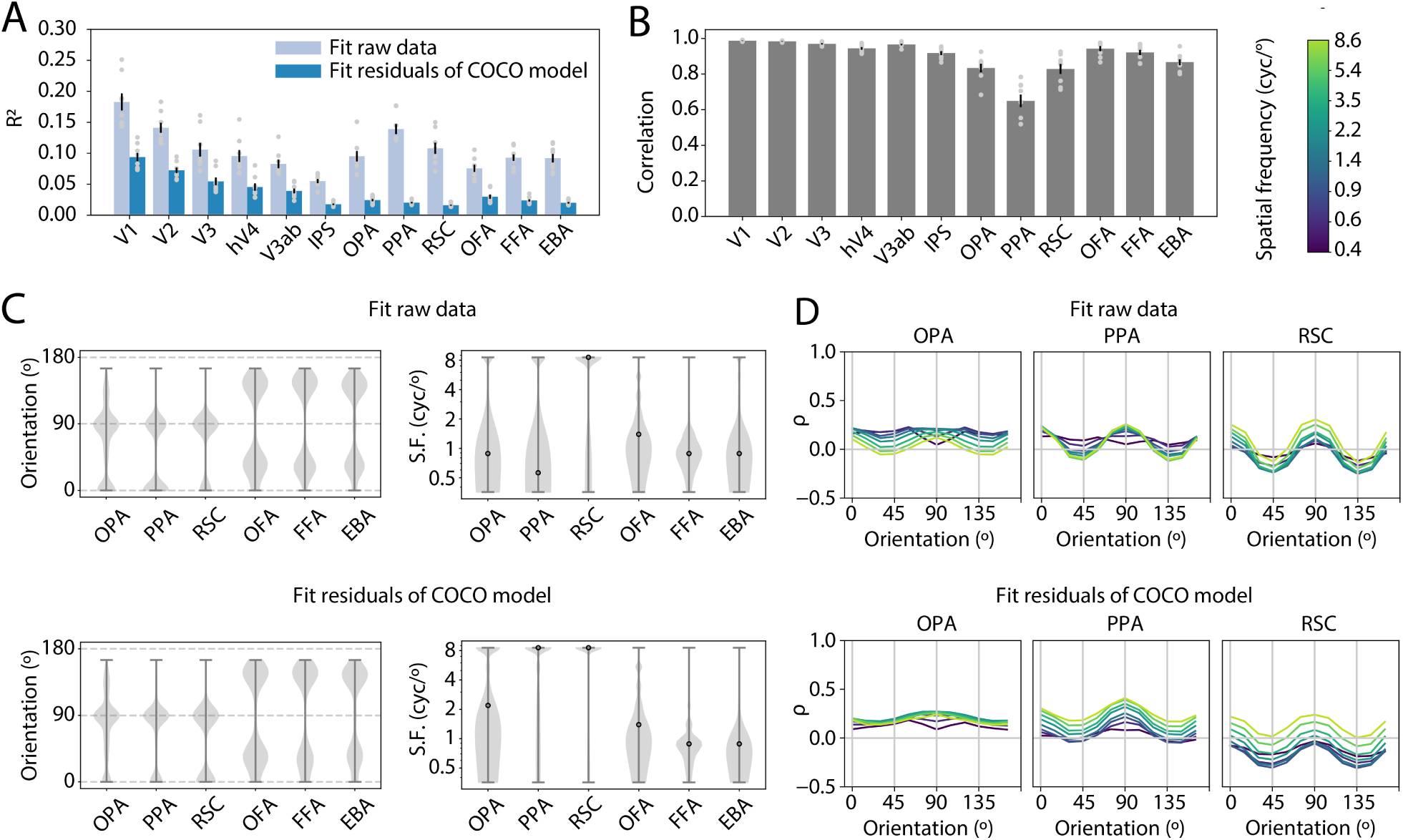
Visual feature selectivity persists after regressing out the contributions of explicit category selectivity from the voxel responses. **(A)** Overall model performance (R^2^) when fitting the Gabor encoding model using the raw data or the residuals of the COCO-all semantic encoding model (a model whose features describe the presence of various object and stuff categories in the image; see *Methods: Semantic Features* and *Methods: Regressing Out Category Selectivity* for details). **(B)** The average correlation between individual voxel feature sensitivity profiles (i.e., sensitivity for each of the 96 Gabor features) that were estimated using the raw data or the residuals of the COCO-all semantic model, averaged across voxels in each ROI. High values indicate that feature selectivity was similar whether fitting on the raw data or the residuals. In **(A)** and **(B)**, bar heights and error bars indicate mean ± 1 SEM across 8 participants, gray dots indicate individual participants. **(C)** The distribution of preferred orientation (left panels) and preferred spatial frequency (right panels), when fitting using either the raw data (top panels) or the COCO-all residuals (bottom panels). Note that the top panels are similar to Figure 3C-D, except that the voxels have been additionally thresholded based on R^2^ for the Gabor model fit to the COCO-all residuals (see Supplementary Table 4 for voxel counts after thresholding). **(D)** Feature sensitivity profiles as a function of orientation, averaged across voxels in several example ROIs (each colored line is a different spatial frequency level, colorbar in top right). The top three panels represent feature sensitivity estimated from the raw data, while the bottom three panels represent feature sensitivity estimated from the COCO-all residuals.

Visual feature selectivity was also consistent when fitting the encoding model on images from only one category at a time (Figure 6; see *Methods: Fitting Models for Single Categories* for details). Similar to the previous analysis, we compared the feature sensitivity profiles for models fit on images of different categories, and found that correlations were positive on average in all areas, with the highest values in early visual cortex and OFA (Figure 6A). The correlation was lowest in place-selective ROIs, followed by IPS and EBA. The sensitivity of feature selectivity to category appeared to depend on which specific categories were compared: for instance, in RSC, sensitivity to category was highest (i.e., correlation was lowest) when comparing between models fit on outdoor images versus indoor images, but in EBA, sensitivity was highest when comparing models fit on large versus small images. To investigate these effects further, we visualized the distribution of peak orientations and peak spatial frequencies, as well as the orientation and spatial frequency sensitivity profiles, across models fit to images of one category at a time (Figures 6B-D, Supplementary Figures 7, 8). For comparison, we also generated a sub-sampled set of images for each semantic dimension that was evenly balanced with respect to the two categories of interest; the number of images used for fitting was the same for this image set and the two single-category image sets. In PPA, the distribution of orientation peaks was concentrated at horizontal (90°) orientations when fit on outdoor images, but concentrated at vertical (0°) orientations when fit on indoor images. In spatial frequency, more PPA voxels appeared to prefer high spatial frequencies when fit on either indoor only or outdoor only, as compared to the balanced set (Figure 6B). Additionally, the feature sensitivity profiles in PPA exhibited an overall upward shift for outdoor images over indoor images – potentially suggesting an overall increase in sensitivity for the Gabor model features when measured on outdoor images. In EBA, a shift in the distribution of peak orientations was observed when fitting on large images versus small images, in that when fitting on large images or a balanced set of large and small images, most voxels exhibited a preference for diagonal orientations, while fitting on small images resulted in a group of voxels with peak sensitivity for horizontal (Figure 6D). As in PPA, an overall vertical shift of the sensitivity profiles was observed across categories, with EBA tending to have higher sensitivity values on average when fit with large images as opposed to small. Finally, FFA exhibited stable patterns of orientation and spatial frequency sensitivity when comparing between model fits on animate images, inanimate images, or a balanced set (Figure 6C), suggesting that feature sensitivity in FFA was not dependent on animacy coding.

**Figure 6:**
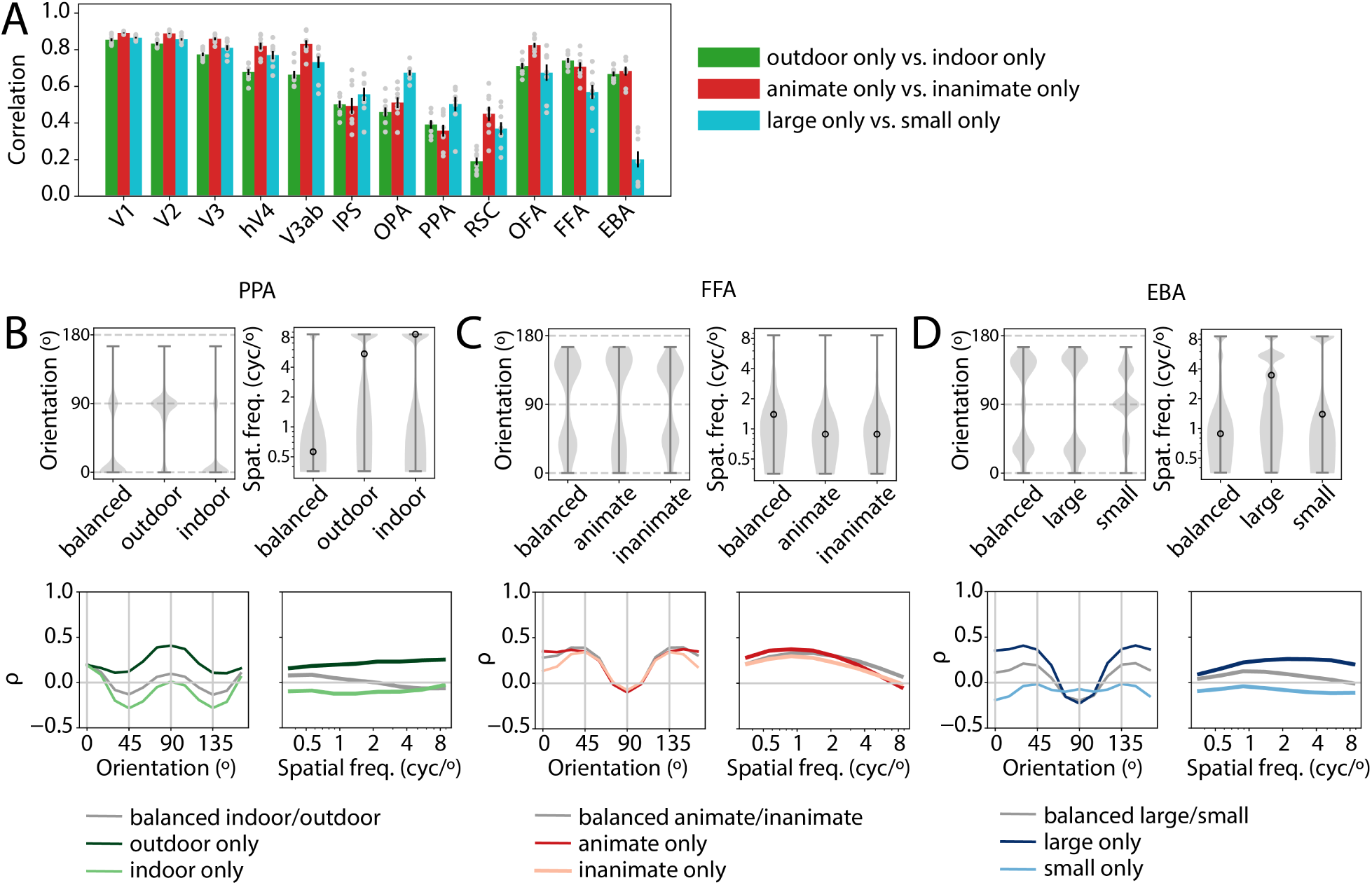
Visual feature selectivity is similar across semantic categories, with interactions evident in some areas. **(A)** The correlation between feature sensitivity profiles for individual voxels estimated using images from just one semantic category at a time (see *Methods: Fitting Models for Single Categories* for details). Error bars indicate mean ± 1 SEM across 8 participants, gray dots indicate individual participants. **(B)** Feature sensitivity for voxels in PPA, comparing between models that were fit using either only outdoor images, only indoor images, or a balanced set of indoor and outdoor images. Top panels show the distribution of preferred orientations and preferred spatial frequencies across voxels, bottom panels show orientation and spatial frequency sensitivity profiles, averaged over all voxels in PPA. **(C)** and **(D)** are analogous to **(B)**, but, respectively, show FFA sensitivity compared between animate/inanimate images, and EBA sensitivity compared between large/small images. For the distribution of feature selectivity across all category-selective ROIs for each individual category, see Supplementary Figures 7 and 8.

## Discussion

To extract meaning from the world, intelligent systems leverage regularities such as the co-occurrence statistics between sensory features and semantic categories. Under this view, the high-level organization of semantic representations in the human brain should reflect natural scene statistics – a hypothesis we investigated by examining the relationship between low-level visual and high-level category selectivity in visual cortex. First, we demonstrated that semantic dimensions are associated with distinct signatures of low-level visual features, and that information about these distinctions is distributed non-uniformly throughout the visual field. Second, we compared the selectivity for these same features across brain areas thought to play different roles in high-level visual processing, and found meaningful correspondences. Cortical regions tended to over-represent components of the visual world that were informative for detecting their preferred category. Together these results indicate that low-level feature biases observed throughout visual cortex reflect the structure of the visual environment, and thus may provide organizational scaffolding for high-level category representations.

### Biases in low-level feature selectivity

With respect to the dimension of orientation preference, our results revealed that place-selective visual ROIs and IPS were preferentially selective for the cardinal orientations (0°/90°), while early visual, face- and body-selective ROIs exhibited stronger selectivity for diagonal orientations. The finding of cardinal bias in place-selective areas is consistent with past reports [21], but the lack of such a bias in early visual areas is surprising in light of past work demonstrating that at the single-unit level, more neurons in V1 and other early areas tend to be tuned for vertical and horizontal orientations than diagonals [43, 44, 45]. A possible explanation for this discrepancy is the fact that we measured neural responses to natural image stimuli, whereas most past studies have used simple, synthetic stimuli such as oriented gratings. Indeed, at the behavioral level, it has been suggested that the commonly-observed “oblique effect”, in which observers tend to show better performance close to the cardinal orientations and worse at the obliques [46], may reverse when measured with more naturalistic, broad-band spatial frequency images [47]. This “horizontal effect” has been explained in terms of a divisive normalization mechanism in which the normalization pool acting to suppress neuronal activity might be larger for cardinal orientations than oblique orientations [47]. Given that horizontal and vertical orientations are overall more common in natural images [48, 49, 50], this mechanism might serve to suppress the most common orientations in natural images while enhancing processing of uncommon (i.e., unexpected) information [47]. Such an explanation would be consistent with the idea that orientation anisotropies in early visual cortex reflect efficient coding of natural image statistics and suppression of redundant information [51, 52, 53]. At the same time, another factor that may contribute to the discrepancy between our results and past electrophysiology work is the difference in recording method. Since the fMRI signal reflects pooled synaptic activity over many neurons, it may give less precise measures of orientation selectivity than single-neuron recordings [54], and could also be differentially sensitive to the divisive normalization mechanism described above. Supporting this, some past fMRI studies, in agreement with our results, have found greater activation of early visual cortex for oblique than cardinal orientations [55, 56]. On the other hand, one fMRI study found that the relative strength of activation for oblique orientations versus cardinals is dependent on stimulus properties such as contrast [57], which lends support to the idea that orientation selectivity may exhibit a different functional signature when measured using natural images versus gratings.

Considering these factors *in toto*, one interpretation of the difference between early visual cortex and scene-selective cortex in our results is that feature selectivity in early areas may be more strongly constrained by generic efficient coding of images, without regard to semantic content, while higher visual areas may reflect stronger semantic constraints, including the association of cardinal orientations with scene-diagnostic content such as large, inanimate objects. An increase in the magnitude of cardinal orientation bias from early to late stages of visual processing is also consistent with past reports of stronger cardinal biases at deeper layers of a convolutional neural network [50]. This pattern is also compatible with our findings in face- and body-selective ROIs, in which orientation biases were similar to those in early areas, but with a slight shift in the distribution of preferred orientations toward vertical (shifting from peaks at 45°/135° to peaks at 30°/150°; Figure 3). Given that 30° and 150° were the orientations most strongly associated with animacy (Figure 2), this shift in orientation biases may reflect an increase in the influence of semantic constraints on feature selectivity when moving from early visual to higher visual regions.

With respect to a second low-level dimension, spatial frequency, we again found evidence for differences in the spatial frequency selectivity of voxels across different ROIs. One notable finding was that RSC and PPA each exhibited a large proportion of voxels tuned for high spatial frequencies, although in PPA this was only the case after regressing out the contributions of category selectivity from the voxels’ responses (Figure 5). The finding of high spatial frequency selectivity in PPA has been reported previously [24, 58], although others have reported relatively less sensitivity to high spatial frequency in RSC [58]. Further supporting the relevance of high spatial frequencies for processing in scene regions, a recent study also showed that decoding of scene categories in PPA and RSC was driven predominantly by high spatial frequency information [59]. Our results are consistent with this finding, and we further show that high spatial frequency features carry information about the distinction between indoor versus outdoor scenes (Figure 2). Thus, the high spatial frequency biases in scene-selective ROIs appear to be consistent with their putative roles in scene processing.

### Biases in spatial selectivity

With respect to spatial selectivity, we found evidence for biased coverage of the visual field in several ROIs, as well as corresponding biases in the distribution of category-diagnostic feature information around the visual field. In particular, category-selective areas OPA, OFA, and EBA all showed bias in favor of the lower half of the visual field (Figure 4), and, correspondingly, the decodability of object properties animacy and real-world size was higher in the lower visual field (Figure 2). These findings are consistent with past reports of high-level visual areas on the lateral surface of the brain tending to be biased in favor of the lower hemifield [60, 61, 42, 62], as well as with past findings of scene-diagnostic objects tending to be more concentrated in the lower hemifield [63]. We build on this work by demonstrating that two key high-level object properties, animacy and real-world size, are more easily decodable from features in the lower versus the upper visual hemifield, and that this spatial bias in the visual environment is reflected in patterns of selectivity within specific visual areas. Of note, there is some evidence from past studies for an upper visual field bias in visual areas on the ventral surface, including PPA and FFA [42, 64], although at least one study found no evidence for an upper visual field bias in FFA [65]. We did not find evidence for a visual field bias in FFA or PPA in our main set of pRF analyses. However, in a supplementary analysis, where we fit pRFs using a different model, some evidence for an upper visual field bias was observed in both PPA and RSC, although not FFA (Supplementary Figure 6). These disparate results suggest that methodological choices may be a contributing factor to the different measurements of visual field bias in ventral visual cortex across studies.

In addition to biased coverage of the lower versus upper visual hemifield, we observed biases along the foveal to peripheral axis. The most foveally biased visual area was OFA, while scene-selective areas PPA and RSC had relatively more voxels with peripheral pRFs (Figure 4). These findings are broadly consistent with past findings of foveal and peripheral biases in face- and scene-selective areas, respectively [26, 27, 65]. We also extended this past work by showing that information carried by Gabor features about object properties animacy and realworld size was higher for more central pRF positions, while information about the indoor versus outdoor scene distinction was higher for more peripheral pRF positions. The role of peripheral and/or global image features in supporting scene perception has been suggested by past work [66, 67, 64]. The correspondence between these sets of findings is consistent with the idea that face-selective areas are biased in favor of the detailed parts of the image that may be informative for face detection (i.e., animacy in the central visual field), while scene-selective areas are biased in favor of the parts of the image more informative for processing large-scale scene distinctions.

### Interactions between feature and category coding

Taken together, the above results provide evidence for reliable, selective sensitivity to low-level features throughout the entire visual cortical hierarchy. Moreover, we demonstrate that the feature selectivity seen in higher visual cortex cannot be attributed solely to coding of semantic category information (Figure 5, Figure 6). That is, even after removing the contributions of explicit category selectivity from voxel responses, a Gabor encoding model predicted a substantial amount of the variance, and biases in feature selectivity for each brain region were largely stable following this manipulation. Our observation is in keeping with past work showing low-level feature selectivity, and biases for particular features, in higher visual cortex regions [68, 69, 24, 21, 19], and supports the interpretation that higher visual cortex responses reflect a mixture of visual and categorical information.

Several theoretical proposals have been put forth to explain the relationship between visual and category selectivity. One hypothesis is that functional selectivity for a category reflects a combination, either additive or nonlinear, of selectivity for a set of underlying features that comprise the category [32]. Under such a coding scheme, neural populations might be selective for sets of features that are diagnostic of a particular category, but also exhibit some residual selectivity for the category itself [33, 70]. Similarly, work using synthetic stimuli has shown that mid-level visual features, in the absence of semantic information, can generate signatures of categorical selectivity in ventral visual cortex, but that other (possibly semantic) features may be required to elicit the strongest level of category selectivity in these regions [71]. Our results do not speak directly to the presence of residual categorical effects, but they do lend support to the hypothesis that visual feature selectivity in high-level cortex regions may play a role in the computation of categories through the representation of category-diagnostic features.

Interestingly, our results also suggest interactions between feature coding and category coding, particularly in place-selective visual regions (Figure 6). In PPA, the relative strength of biases for the two cardinal orientations shifted depending on whether fitting was performed on outdoor or indoor images, with stronger sensitivity (*ρ*) for 90° observed when fitting on outdoor images versus indoor. This pattern may reflect the contributions of several different factors. First, it is possible that differences in the statistical distribution of features between outdoor and indoor images can explain the effect. As shown in our first analysis (Figure 2), outdoor images have more energy at 90° orientations than indoor images, and it is possible that the presence of these outdoor images, dominated by horizontal orientations, is required in order to measure the full extent of sensitivity to horizontal orientations in PPA voxels. Alternatively, it is possible that PPA voxels show overall higher responses to either indoor or outdoor images, which impacts their signal-to-noise ratio in one condition, and thus impacts our detection of feature sensitivity. Analyses addressing this question suggested that PPA may be biased toward indoor images over outdoor (Figure 3E), a finding also reported elsewhere [72]. Based on this bias and the fact that average orientation sensitivity values tended to be higher when estimated from the outdoor images, this may indicate that the overall lower signal for outdoor images produced greater sensitivity for detecting orientation sensitivity (perhaps due to ceiling effects in responses to the indoor images). Similar arguments can be made for the other feature-category interactions, including the apparent change in orientation sensitivity when fitting on large versus small images in EBA. However, our present results do not readily allow us to differentiate these hypotheses, and it is possible that the patterns are driven by a combination of image statistics, signal-to-noise ratio and some form of mixed selectivity for features and categories in higher visual areas. This is one important direction for future work.

### Associations between low-level features and semantic categories

Our measurements of the associations between low-level features and semantic categories (Figure 2) are consistent with several earlier studies. For example, Torralba and Oliva [28] compared the spectral content for a range of image categories and found differences between scene categories, as well as between images of different object classes. Consistent with our findings, certain classes of outdoor images, such as forests and cities, showed larger power at high spatial frequencies. Similarly, certain classes of outdoor images, such as beaches and highways, had higher power at horizontal orientations. At the object category level, Torralba and Oliva reported spectral differences across images containing animals versus other objects, which is consistent with our observation that certain orientation channels were associated with animate over inanimate objects. However, an important difference between our approach and theirs is that we labeled images according to their animacy or inanimacy in a spatially-specific manner – when we computed the features that were correlated with animacy, we only used features extracted from image patches that actually contained either an inanimate object, an animal, or a person. Thus, the features correlated with animacy in our analyses are not likely to have been as strongly driven by background content as those used in Torralba and Oliva’s work. Despite this, there is still a correspondence between our finding that diagonal orientations were positively associated with the animate-inanimate axis, and Torralba and Oliva’s finding that images with animals had relatively more power at diagonal orientations (i.e., were less strongly dominated by cardinal orientations) than images with other types of objects. Our finding that vertical and horizontal orientations were associated with objects having a large real-world size is not entirely predicted from this past study, but may be consistent with their observation of strong cardinal biases in scenes including cars. The relationship between cardinal orientations and large objects is also consistent with the observation that objects having a large real-world size are dominated by boxy contour features [30, 22]. Importantly, our results build on these past observations by providing detailed comparisons of the correspondence between a fine-grained bank of Gabor features and three high-level semantic dimensions. These measurements provide a foundation for interpreting the fine-grained biases in feature representations within visual cortex.

### Conclusion

The structured relationship between category selectivity and visual feature selectivity in the brain may emerge due to multiple factors, including visual experience during development and functional or anatomical neural constraints. One account for the organization of visual cortex suggests that from infancy, the primate brain includes a “proto-organization” that may be a precursor to the large-scale maps of category and feature selectivity observed in adult brains [73, 74]. These early topographic constraints, including selectivity for low-level features such as spatial frequency and curvature, may interact with visual input to constrain where mature category-selective visual areas will develop [75]. For example, retinotopic biases present in the early visual system interact with the portion of the visual field in which certain classes of stimuli tend to fall (i.e., faces and words tend to be foveated, buildings tend to land in the periphery), and this may lead to a correspondence between retinotopy and category selectivity [26, 27]. Weak early biases for features such as curvature may similarly lead cortical populations to develop selectivity for those categories in which such features are prominent [74]. Our results provide some support in favor of this hypothesis, in that we find a correspondence between the features and spatial positions associated with a given category and the low-level tuning of neural populations involved in processing that category. Thus, it is plausible that experience with the statistics of natural images during development, along with some degree of early topographic organization, may be sufficient to predict the organization of feature, spatial, and category selectivity in the adult visual system.

Category-selective visual areas appear to contain representations of diagnostic low-level features, which could suggest that these low-level feature representations play a functional role in category perception. Though our study did not directly assess this functional role, past work supports the idea that low-level visual features may influence behavioral judgments of object and/or scene category. For example, rapidly classifying scenes into basic-level categories may be mediated by global properties associated with the spectral content of scenes [29, 66, 67]. Detecting object categories, such as animals, may also be supported by spectral differences across images [28]. At the same time, other work has suggested that in the case of animal detection, spectral features may not be sufficient to predict behavior [76] and that features more complex than spectral content, such as mid-level textural features [30, 31] or contour junctions [77] may be more useful in computing semantic properties of objects and scenes. Additional experiments are needed to determine whether the feature biases we measured play a functional role in behavior, but our results do add to a growing body of evidence that low-level features contain potentially informative cues to the semantic meaning of images. We build on this past work by showing that these cues are reflected in the brain across a range of visual areas and semantic category distinctions.

Our analyses focused on well-studied category-selective and retinotopic regions in visual cortex in order to facilitate comparisons with past studies. At the same time, the large-scale organization of higher visual cortex may be better understood in terms of broad visual feature dimensions that are mapped onto the cortex in a continuous fashion [32, 78, 79], rather than as a discrete set of regions processing disparate types of information. In this framework, the biases that we observed in feature tuning within functionally-localized regions of visual cortex may be interpreted as reflecting the correspondence between maps encoding features at different levels of complexity. This view is also compatible with our observation that there was heterogeneity in the feature sensitivity profiles measured for voxels within a given region of interest (e.g., the spread of distributions in Figure 3). It seems plausible that the heterogeneity in feature tuning of voxels across an ROI is related to heterogeneity in their semantic selectivity; again, future work will be needed to explore this possibility.

Overall, our results provide evidence in favor of the theory that representations of low-level features within category-selective regions of visual cortex are aligned with the high-level computational goals of these regions. Moreover, this principle appears to hold for neural populations across a wide range of visual areas and for multiple semantic dimensions. These findings suggest that the computation of semantic meaning in the visual system may reflect contributions from features at multiple levels of complexity.

## Methods

### Human Participants and Acquisition of fMRI Data

We used a large-scale publicly available dataset, the Natural Scenes Dataset (NSD), for all analyses. A detailed description of the data is provided in [37]. Briefly, the NSD includes measurements of whole-brain BOLD fMRI from 8 participants who each viewed between 9,000-10,000 colored natural scenes over the course of 30-40 scan sessions. All functional scans were conducted at 7T using whole-brain gradient-echo EPI at 1.8 mm resolution and 1.6 second repetition time. Images were taken from the Microsoft Common Objects in Context (COCO) database [36], and were square-cropped and presented in color, at a size of 8.4° x 8.4° (° = degrees of visual angle). Of the approximately 10,000 images viewed by each participant, ∼9,000 images were seen only by that participant, and ∼1,000 were overlapping across participants. Each image was viewed for a duration of 3 seconds, with 1 second between trials. Over the course of the experiment, each image was viewed approximately three times, for a total of roughly 30,000 trials per participant. Participants were required to fixate centrally on a small fixation dot superimposed on each image, while performing a task in which they reported whether or not each image had been presented before in any session.

### Pre-processing of fMRI Data

fMRI data were preprocessed by performing one temporal interpolation to correct for slice time differences and one spatial interpolation to correct for head motion within and across scan sessions [37], resulting in volumetric fMRI time-series data at 1.8 mm resolution in subject native space. Beta weights for each voxel and each trial were then estimated using a general linear model. To improve the signal-to-noise ratio in estimating beta weights, a three-stage procedure was used which consisted of selecting a hemodynamic response function (HRF) from a library of candidate HRFs, denoising the data using a set of noise regressors estimated from voxels not related to the experimental paradigm, and using fractional ridge regression to regularize the beta weight estimation on a single voxel basis [80]. Finally, beta weights for each voxel were averaged over trials on which the same image was shown (typically three trials per image), and the averaged beta weights for each image were used for all further analyses.

### Defining Regions of Interest (ROIs)

Each NSD participant also performed several runs of a category functional localizer task [81] and a population receptive field (pRF) mapping task [82]. The pRF mapping task was used to define early retinotopic visual ROIs V1, V2, V3, and hV4 [37]. The category localizer task was used to define face-selective ROIs (occipital face area/“OFA” and fusiform face area/“FFA”; for our analyses we combined FFA-1 and FFA-2 into a single FFA region), place-selective regions (parahippocampal place area/“PPA”, occipital place area/“OPA”, and retrosplenial cortex/“RSC”), and a body-selective region (extrastriate body area/“EBA”). In addition to these functional ROIs, we also utilized a probabilistic atlas [83] to define areas V3ab and the IPS (intraparietal sulcus; all six sub-regions of IPS0-5 were combined into a single region). Since the probabilistic atlas also included definitions for early visual areas V1, V2, V3, and hV4, which were already defined using the pRF mapping task, we combined both sets of definitions for these regions, and where they disagreed we deferred to the pRF-based definitions. For example, if a voxel was labeled as V1 in the atlas and was not included in the pRF-based labels, it was added to V1, but if it was labeled as V1 in the atlas and V2 in the pRF-based labels, it was kept as V2. As a result, the retinotopic ROI labels were non-overlapping with one another. To prevent the category-selective ROIs from overlapping with the retinotopic ROIs, we removed any voxels that overlapped from the retinotopic ROIs and included them only in the category-selective ROI to which they corresponded; the area most affected by this was V3ab, which was originally overlapping with OPA. To prevent category-selective ROIs from overlapping with one another (which can happen for regions defined based on different contrasts), we removed the overlap by prioritizing face-selective definitions over body and place-selective definitions, and prioritizing place-selective definitions over body-selective definitions. Thus, the final set of 12 ROIs were entirely non-overlapping. Finally, we applied an additional threshold to the voxels in each ROI based on their noise ceiling (i.e., the theoretical proportion of variance in the data that can be predicted; for details on noise ceiling calculation see [37]), using a threshold of 0.01. A full list of ROI sizes, following noise ceiling thresholding, is presented in Supplementary Table 1).

### Encoding Model Fitting

#### Overview

We fit an encoding model for each individual fMRI voxel which captured its spatial selectivity as well as its feature selectivity [84] (Figure 1B). Fitting was done for two different visual feature spaces and a semantic category feature space (see *Methods: Feature Spaces*). The basic procedure was to first loop over a grid of candidate population receptive fields (pRFs, see *Methods: Population Receptive Fields (pRFs)*) for each voxel, and use regularized regression to fit the weights of a linear model that describes the voxel response as a weighted sum of the feature activations corresponding to that pRF. The best-fitting candidate pRF and the feature weights for that pRF made up the final encoding model. We then evaluated that model’s ability to predict voxel responses on a held-out partition of data. The fitted encoding models were also used to estimate voxel feature selectivity. Below, we outline each of these steps in detail.

#### Population Receptive Fields (pRFs)

Following from past work [38, 40, 85, 84], we modeled the spatial selectivity of each fMRI voxel using a 2-dimensional Gaussian response profile over the spatial extent of the viewed image. This approach is similar to classic approaches of fitting spatial receptive fields for single neurons in visual cortex, and is often termed a population receptive field (“pRF”) to denote summation over the population of neurons in a voxel. The Prf can be described by three parameters, *x*_*0*_, *y*_*0*_, and *σ*, where [*x*_*0*_, *y*_*0*_] and *σ*, respectively, indicate the center and standard deviation of the 2-dimensional Gaussian response profile:

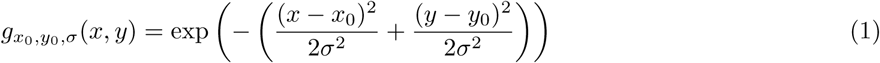

To select each voxel’s optimal pRF, we constructed a grid over candidate pRF parameters. Our grid had even spacing between adjacent pRF centers in terms of their polar angle position (*θ*), and nonlinear spacing in eccentricity (*r*), where candidate centers were more closely spaced closer to the center of the visual field. The purpose of the nonlinear eccentricity spacing was to account for the cortical magnification factor in human visual cortex, where the neuronal sampling of visual space is more dense close to the fovea [86]. More concretely, our 16 candidate polar angle positions were linearly spaced, ranging from 0-337.5° in steps of 22.5°, and our 10 candidate eccentricities were logarithmically spaced, ranging from 0° - 7°. Our 10 *σ* values were also spaced logarithmically, with *σ* ranging from 0.17° to 8.4° (note that throughout this paper, we use ° to refer both to geometric angle, in the context of polar angle and orientation, and degrees of visual angle, in the context of eccentricity and size of pRFs). To generate the complete grid, we first computed every possible combination of *r, θ*, and *σ*, which resulted in 1600 pRFs. We then converted the centers from polar angle coordinates [*r, θ*] into Euclidean coordinates [*x*_*0*_, *y*_*0*_]. Finally, we eliminated any pRFs that landed completely outside the image region (an 8.4°x8.4° square), by the criteria that their rough spatial extent (center±*σ*) was non-overlapping with the image region. The result of this was that for the most peripheral pRFs, only the larger size values were included in the grid. The final grid included 1456 pRFs in total.

#### Encoding Model Design

Our encoding model framework assumed that each voxel response could be modeled as a linear weighted sum of the activations in a set of underlying feature channels. In mathematical notation, this can be formulated as:

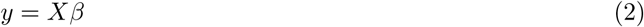

Where *y* is a column vector of length *n* containing the voxel responses across *n* images, *β* is a column vector of length *w* containing *w* weights, corresponding to each of (*w* − *1*) feature channels plus an intercept. *X* is the design matrix for the feature space, describing the activation in each feature channel for each image plus a column of ones, size [*n* × *w*].

The features in each of our feature spaces were computed in a spatially specific manner [84], such that the design matrix depended on the pRF parameters *x*_*0*_, *y*_*0*_ and *σ*. Referring back to the pRF definition given in Equation (1), the design matrix can be expressed as:

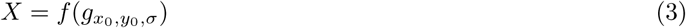

Where *f* is a function that depends on the feature space under consideration. In most cases, *f* simply refers to taking the dot product of the pRF with a spatial feature map describing the activation of some feature channel at each position in the image. In those cases, for the spatial activation map *a*_*t,c*_ (size *p* × *p*) corresponding to image *t* and feature channel *c*, and pRF profile *g* having the same resolution as *a*, the corresponding element of *X* can be computed by:

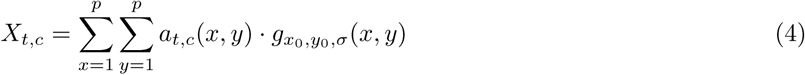

But this form differs when we use a semantic category feature space; see *Methods: Semantic Features* for details.

Note that Equation (4) requires generating each pRF with a flexible resolution to fit the resolution of the feature map output from a given model. To achieve this, we scaled the *x*_*0*_, *y*_*0*_ and *σ* parameters along with the feature map resolution, so that they always corresponded to the same positions in image coordinates.

Also note that when *f* takes the form described in Equation (4), the *σ* parameter of the pRF does not provide a complete description of the spatial extent of the image that contributes to computing *X*. This is because each pixel in the activation map *a* has its own pooling region, determined by the operations used to compute that activation (e.g., the kernel size of a convolution). Thus, *σ* should be interpreted as a lower bound on the pRF size rather than an exact estimate of its size [84].

#### Model Fitting Procedure

Following previous work [87, 88, 89], we solved for the weights for each voxel using ridge regression (L2-regularization). The ridge regression estimator of *β* is given by:

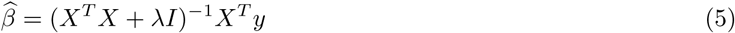

Where *I* is a [*w* × *w*] identity matrix, *λ* is a regularization parameter, and *y* is a vector containing the voxel response for *n* images. Once 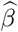 is computed, the voxel response can be predicted from the design matrix associated with any arbitrary stimulus input, by:

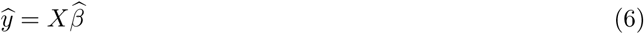

The regularization parameter (*λ*) was selected using cross-validation on a per-voxel basis, from a set of 10 candidate *λ* values logarithmically spaced between 0 - 10^5^. The full cross validation procedure is detailed below (see also Figure 1B).

First, we held out ∼1,000 images from the ∼10,000 total images for each participant, to serve as a validation set. The validation set images always consisted of the set of “shared images” that were seen by all participants (see *Methods: Human Participants and Acquisition of fMRI Data*). The remaining ∼9,000 images made up the training data. Before fitting, we z-scored the values in each column of the design matrix, separately for the training images and validation images. To select the ridge parameter and the best pRF parameters for each voxel, we held out a random 10% of the training data as a nested validation set. We then used the remaining 90% of training images to compute 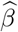 for each of our candidate *λ* values and candidate pRF models (recall that the pRF parameters determine the design matrix *X* used, see Equation (3)). For each of the candidate pRFs and *λ* values, we computed a prediction of the nested validation data 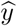 based on the estimate of 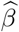, and computed the loss of that estimate, 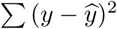. The pRF parameters and *λ* value that resulted in the lowest loss were selected as a the best pRF parameters and *λ* for that voxel. Correspondingly, the 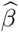 associated with that pRF and *λ* were selected as the best weights for that voxel. Finally, these best fit parameters were used to predict each voxel’s response on the held out validation data, and we computed the coefficient of determination (*R*^2^) between *y* and 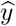 as a measure of overall model accuracy.

The above procedure describes our method for fitting spatial and feature selectivity simultaneously, as done in [84]. However, once a stable estimate of the pRF for each voxel has been obtained using some feature space, we can adapt this method to fit just the feature selectivity for each voxel (for any arbitrary feature space), assuming that its spatial selectivity remains fixed. To do this, we use each voxel’s pre-computed pRF estimate to select the correct design matrix *X*, and fit its 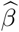 for that design matrix only, for each candidate *λ* value. We follow the same procedure as outlined above with respect to training/validation image splits and *λ* selection.

For all analyses presented here, our approach was to first fit both the feature and spatial selectivity of each voxel using the AlexNet concatenated feature space (see *Methods: AlexNet Features*). Once the pRF estimates were obtained based on AlexNet, we assumed that these pRFs remained fixed for all other feature spaces and used them to fit each voxel’s weights for all other feature spaces of interest (i.e., Gabor and COCO-all; see *Methods: Feature Spaces*). We chose the AlexNet feature space [39] for the first pRF fitting step because AlexNet has often been used to model voxel responses in human and non-human primate visual cortex, and gives good predictive performance across a range of cortical areas [90, 91, 92, 23]. In our data, AlexNet generally yielded higher predictive accuracy than the Gabor model in higher-level visual areas (Figure 3A), which meant we had more voxels in those areas whose R^2^ met the criterion for inclusion in our pRF analyses (Supplementary Table 2, Supplementary Table 3). Similar results for both spatial and feature selectivity were obtained when we fit the entire model, including pRFs, from scratch on the Gabor feature space (Supplementary Figure 6).

### Feature Spaces

#### Overview

For each stimulus image viewed by fMRI participants, we extracted several different sets of features that were intended to capture different aspects of the image’s visual and semantic content. These features were used to construct voxelwise encoding models, as well as to estimate the statistical associations between lower-level features and semantic categories (see *Methods: Measuring Feature-Semantic Associations*). Each set of features was extracted in a spatially-specific manner, such that the features associated with each pRF grid position described the visual or semantic content within a specified region of the image only. Unless otherwise specified, all feature extraction was performed on grayscale images at a resolution of 240 × 240 pixels. Below, each feature space is described in detail.

#### Gabor Features

Our first set of features was based on Gabor filters that extract the energy at specified orientations and spatial frequencies (see Figure 1A). Similar models have previously been used to model the responses of early visual cortex to natural images [84, 93, 94]. Our Gabor filter bank included filters at 12 unique orientations, linearly spaced between 0-165° in increments of 15°. Each filter consists of a 2-dimensional complex-valued sinusoid with a specified frequency and orientation, multiplied by a 2-dimensional Gaussian envelope. The real and imaginary components of the sinusoid are at 0° and 90° phase, respectively, to form a quadrature pair. The final activation of each filter was obtained by convolving both the real and imaginary filters with the input image, squaring the output of both the real and imaginary parts, summing the real and imaginary parts, and taking the square root. We then applied a nonlinearity to the resulting activation values, 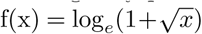.

We applied this bank of filters at 8 spatial frequencies that were logarithmically spaced between 0.35 and 8.56 cycles per degree of visual angle (cyc/°). To achieve filtering at each frequency, we first resized the input images to an appropriate size (i.e., smaller for lower frequencies) using bilinear resampling. We then applied the stack of filters, which were always a fixed size and frequency in pixels (12×12 pixels and 4.13 pix/cycle), to the resized images. The end result was a set of 8 stacks of feature maps, one for each spatial frequency, where the height and width dimensions of each stack depended on its corresponding spatial frequency. Each stack contained 12 feature maps for the 12 orientation channels.

Finally, to extract the feature activations within each pRF in our grid, we took the dot product of the pRF with each feature map to obtain a single value for the activation in each feature channel (Equation (4)). This resulted in a 96-dimensional feature space, computed separately at each pRF grid position.

#### AlexNet Features

We extracted visual features from a convolutional neural network model referred to as “AlexNet”, which is trained on a 1000-way image classification task (see [39] for more details on the model’s construction and training). We extracted activations from the first five convolutional layers of a pre-trained AlexNet model. Activations were extracted after the rectifying nonlinearity (ReLU) function that follows each convolutional operation. To extract the features for each pRF grid position, we took the dot product of each feature map with the pRF of interest (Equation (4)), which resulted in a single value for each feature channel. The dimensions of the resulting feature sets corresponding to each AlexNet layer are [64, 192, 384, 256, 256].

Before using these features in our encoding models, we reduced the dimensionality of features from each AlexNet layer using principal components analysis (PCA). PCA was always performed using the training data only to solve for the principal components, and then using those components to transform all data into the same subspace. We retained a sufficient number of components to explain at least 95% of the variance in the training data. We performed PCA on the features from one pRF at a time. As a result, the dimensionality of features from different pRFs was allowed to differ following PCA, even though the dimensionality of the features from each pRF was the same before PCA. After performing PCA on the features from each layer, we concatenated the features across all layers.

#### Semantic Features

In addition to modeling the visual features present in our image set, we created a feature set that explicitly modeled the semantic categories present at each spatial position (i.e., each pRF) in each image. To achieve this we took advantage of the object segmentations that are associated with each image in the Microsoft COCO database [36], as well as an additional set of semantic segmentations that label the “stuff” (meaningful but amorphous regions such as sky, walls, etc.) in each image (COCO-stuff; [95]). Each object or stuff instance in each of these labeling schemes is accompanied by a polygon which defines the spatial extent of the instance. To determine whether each instance was overlapping with a given pRF in a given image, we first generated a binary mask for the label polygon, at a resolution of 425 × 425 pixels. We then created a second binary mask of the same size, which captured a circular region ± 2*σ* from the pRF center (see Equation (1)). The pRF was considered to be overlapping with the label if the two masks overlapped by at least 10 pixels. This very lenient overlap threshold was meant to account for the possibility of noise in our pRF parameter estimates, as well as the fact that receptive fields in category-selective regions of visual cortex tend to be large. In initial tests, using a more stringent overlap threshold led to poorer fits of the semantic model.

Using this method, we extracted several sets of semantic features for each image and each pRF. The first set of semantic features, which we termed “COCO-all”, included 80 basic-level object categories, 12 super-ordinate object categories, 92 basic-level stuff categories, and 16 super-ordinate stuff categories, for a total of 200 features. Each feature is a binary label that denotes whether a category is present within the pRF of interest. When building encoding models from this feature set (COCO-all model), we directly used the binary features for each pRF as our design matrix (see Equation (2)). These encoding models were used in order to regress out the contributions of semantic selectivity from voxel responses (see *Methods: Regressing Out Category Selectivity*).

Next, we created three additional semantic feature dimensions that captured coarse-level semantic information about the image contents: indoor-outdoor, animate-inanimate, and real-world-size. These higher-level semantic labels were defined on the basis of the “things” and “stuff” category labels. For example, if a Prf contained any animate object (i.e., person, cat), it was labeled as having the “animate” label, and if it contained any inanimate object, it was labeled with the “inanimate” label. This allows for the possibility that pRFs could be labeled with neither the animate nor the inanimate label, or could be labeled with both. Similarly, the real-world-size label was assigned to pRFs based on whether they contained items we defined as being “small” (e.g., a banana), “medium” (e.g., a stop sign), or “large” (e.g., an elephant).

For the indoor-outdoor distinction, since the indoor or outdoor category is most naturally understood as a property of entire scenes, rather than something which can vary across the spatial extent of a single image, we created labels for the entire image rather than for each pRF separately. To define whether an image was indoor or outdoor, we looked for the presence of particular diagnostic objects or stuff classes (i.e., “car” and “grass” are diagnostic of outdoor images, “bed” and “carpet” are diagnostic of indoor images). When images included both indoor and outdoor classes, we resolved the tie by counting the number of total indoor categories (objects and stuff) and outdoor categories that were included in the image, and taking the maximum. This left only around 7% of images that could not be unambiguously labeled by our method.

When using these high-level labels to compute the semantic associations of Gabor features (see *Methods: Measuring Feature-Semantic Associations*), we reduced each semantic dimension down to a single column which could take only one of two values. For real-world-size, this required excluding the “medium” size category. For all three axes, this required excluding any images whose category membership was ambiguous, due to either both categories being present (e.g., both “animate” and “inanimate”) or neither category being present. The analysis of feature-semantic associations was performed using the remaining images.

### Measuring Feature-Semantic Associations

To investigate the association between each Gabor feature channel and each of our high-level semantic dimensions (indoor-outdoor, animate-inanimate, real-world size; see *Methods: Semantic Features*), we used two approaches. First, we computed a partial correlation coefficient between each of the 96 Gabor features and each binary semantic label of interest, controlling for the contributions of the two other semantic labels. Partial correlations were computed using the linear regression method, which consisted of learning a multiple linear regression that maps from the two “other” semantic dimensions to the semantic dimension of current interest, and another regression that maps from the two “other” semantic dimensions to the Gabor feature of interest. The partial correlation was then obtained by computing the correlation coefficient of the residuals of these two regression fits. This was done for each feature within each pRF individually.

Second, we used a linear decoder to measure the discriminability of semantic categories within the 96-dimensional space spanned by all the Gabor features in each pRF. In contrast to the above method, which focuses on one Gabor feature at a time, this analysis measures how much semantic information can be extracted from the combined pattern of activation across features. We performed decoding using a linear discriminant analysis classifier, implemented in scikit-learn, and a 10-fold cross-validation procedure. To measure decoding performance, we computed d’ from signal detection theory, based on the formula d’ = Z(hit rate) - Z(false positive rate). The hit rate is defined as the proportion of test samples in Condition X accurately classified as belonging to Condition X, and the false positive rate is the proportion of test samples in Condition Y inaccurately classified as belonging to Condition X, and Z is the inverse of the cumulative distribution of the Gaussian distribution. To ensure that comparisons of decoding performance across pRFs were fair, we always used the same number of images to train and test the classifier in each pRF (since semantic labels were created for each pRF individually, the number of images that were unambiguously labeled with each category distinction varied across pRFs). To achieve this, for each semantic dimension, we identified the minimum number of images (*n*) in either category for any pRF. When performing decoding within each pRF, we created a randomly downsampled set of images which consisted of *n* images of each category. This also ensured that the classifier was trained on a perfectly balanced dataset. Note that since this image number matching was done within each semantic dimension separately, the absolute value of decoding performance is not necessarily comparable across semantic dimensions.

To test whether decoding performance varied significantly based on pRF parameters, we estimated the slope of a linear regression fit where the y-coordinate is d’ and the x-coordinate is each pRF parameter of interest (pRF *σ*, eccentricity, horizontal position, and vertical position). Before performing this test, we first removed the most peripheral pRFs (those with eccentricity *>* 4°), and used just the remaining 1,280 pRFs. This was done because in our pRF parameter grid, an even sampling of each *σ*/eccentricity combination was only achievable up to around 4° (see *Methods: Population Receptive Fields (pRFs)*), so removing the more peripheral pRFs ensured that all analyses were performed on a set of pRFs that evenly sampled all combinations of eccentricity and *σ*. To determine whether the relationship with each parameter was significantly different than zero, we used a permutation test where we randomly shuffled the x-coordinate data 10,000 times and re-fit the line. We then computed a 2-tailed *p*-value by computing the number of iterations on which the real slope exceeded the shuffled slope, and the number of iterations on which the shuffled slope exceeded the real slope, taking the minimum and multiplying by 2. The resulting *p*-values were FDR corrected [96] across all pRF parameters and semantic dimensions (12 total values) at *α*=0.01.

Importantly, both of these analyses were performed on a completely independent, randomly selected set of 10,000 COCO images that were *not* seen by any participant in NSD. This was done to ensure that the comparison between our image statistics analyses and our voxel selectivity analyses was non-circular and was not driven by the possibility of overlap in the images. We pre-processed these independent COCO images in a similar manner to the pre-processing used for the NSD images, but instead of computing the “semantic loss” described in [37] to determine cropping boxes, we simply cropped the long edge of each rectangular image in a symmetric way to generate square images.

### Estimating Voxel Semantic Selectivity

We estimated the selectivity of voxel activations for each of our three high-level semantic dimensions (indoor-outdoor, animate-inanimate, real-world size; Figure 3E). Selectivity for each dimension was estimated using a partial correlation coefficient which controls for the contributions of the other two semantic dimensions, analogous to the method described in the previous section (*Methods: Measuring Feature-Semantic Associations*). As in the previous section, each semantic dimension was represented as a binary variable, and any images where the semantic label of interest was ambiguous were not included. Voxel semantic selectivity was always estimated using voxel responses for the validation set images only (see *Methods: Model Fitting Procedure* for how validation set was defined).

### Estimating Voxel Visual Feature Selectivity

After fitting Gabor encoding models to each voxel, we used the fitted models to estimate voxel selectivity for individual Gabor feature channels. These analyses were only performed on well-fit voxels, defined as those whose validation set accuracy, for the Gabor feature encoding model, exceeded a criterion R^2^ value of 0.01 (see Supplementary Table 2 for the number of voxels passing this threshold). To estimate feature selectivity, we first computed the sensitivity of the voxel response (as predicted by the encoding model) to changes in the activation within each feature channel. In mathematical notation, for a given feature channel *c* and a voxel *v* with a fitted encoding model for the feature space of interest, our measure of feature sensitivity *ρ*_*v,c*_ is:

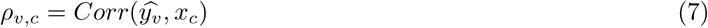

Where 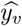 denotes the model-predicted response of the voxel to the validation set images, and *x*_*c*_ denotes the activation in channel *c* for the same validation set images.

This sensitivity measure is meant to capture approximately how strongly each feature channel is “weighted” in the overall encoding model prediction for each voxel. Importantly, however, it is not the same as simply using the raw 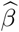 weights as a measure of feature sensitivity. The raw weights of our encoding models are not likely to provide a stable estimate of voxel feature tuning, due in part to the high degree of collinearity between feature channels and in part to the fact that we fit the models with ridge regression, which can lead to biased weight estimates [97, 98]. In contrast, because *ρ*_*v,c*_ is measured using actual validation set data, it provides a measure of the functional alignment between the feature channel and the response of the encoding model, within the context of the real covariance structure of the data. At the same time, we note that *ρ*_*v,c*_ is not intended to dissociate the contributions to the encoding model prediction made by feature channels that are highly correlated.

Once the feature sensitivity (*ρ*) values are computed for each Gabor feature (12 orientations x 8 spatial frequencies) and each voxel, it is straightforward to create a sensitivity profile that captures each voxel’s sensitivity to changes in feature intensity at different positions along the orientation or spatial frequency axis (i.e., Figure 3B). Orientation sensitivity profiles are obtained by plotting the sensitivity values as a function of feature orientation, and spatial frequency sensitivity profiles are obtained by plotting the sensitivity values as a function of feature spatial frequency. Averaging these profiles across values of the other dimension (for instance, averaging orientation sensitivity profiles across all spatial frequency levels), yields an average feature sensitivity profile. These average feature sensitivity profiles were used to compute each voxel’s “preferred” orientation and spatial frequency (i.e., values plotted in Figure 3C-D), through an argmax operation.

### Counting Peaks in Feature Sensitivity Profiles

For each voxel’s average orientation and spatial frequency sensitivity profiles (see previous section), we identified the approximate number of peaks in the curve, as a supplementary analysis to ensure that our results did not depend on assuming a single peak (Supplementary Figure 3). To identify peaks, we first identified all the local maxima in the curve, by comparing each sensitivity value against the values to its left and right. For orientation, we accounted for the circularity of the feature space by wrapping the curve circularly when computing local maxima, while for spatial frequency, we treated the end-points as local maxima if they exceeded the points to their left or right. Next, we removed any peaks that had a negative sensitivity value, since those peaks do not indicate a positive feature preference. Then, we computed the height of each peak by subtracting the minimum value across the sensitivity profile, and divided the height of each peak by the height of the largest peak. Based on the resulting ratio values, we retained only those peaks whose value exceeded 0.50. This ensured that when multiple peaks were counted for a given voxel, each of these peaks was comparable in height.

Based on the finding that many voxels had two or three peaks in their orientation sensitivity profiles, we analyzed the orientation preferences of these multi-peaked voxels by sorting them into groups. For each of the two-peaked (bimodal) voxels, we identified the two orientations at which the two peaks occurred, without regard to the relative height of the two peaks. We then grouped together voxels that were selective for the same pair of orientations. To create the plots in Supplementary Figure 3B-C, we selected the three most common pairs of orientations across all voxels, and combined all other pairs of orientations into a separate group. We used an analogous procedure to group the three-peaked voxels. We did not perform this procedure for spatial frequency, since most voxels had only one true peak in their frequency sensitivity profiles.

### pRF Coverage Analysis

To quantify the visual field biases in each ROI, we computed an aggregated estimate of visual field coverage across all pRFs (similar to [42]). Specifically, we combined all pRF estimates across all voxels in a given brain region (after first thresholding the voxels based on performance of the AlexNet encoding model at an R^2^ value of 0.01, see Supplementary Table 3 for the number of voxels passing this threshold). We combined voxels using an averaging operation across all pRFs, as this takes into account differences in the density of pRF coverage across the visual field, however, similar results were obtained when aggregating using a maximum operation. This led to one aggregated pRF coverage map for each participant and each ROI. We then took the average of these maps in each visual field quadrant to yield an estimate of the coverage for each quadrant.

Next, we analyzed the differences in coverage among ROIs and quadrants by performing a three-way repeated measures ANOVA, with ROI, vertical position, and horizontal position as factors (implemented using the Python package statsmodels). Because this test revealed a significant interaction between ROI and vertical position, we performed *post-hoc* pairwise tests to identify ROIs where there was a difference in coverage between the upper and lower visual fields. Pairwise tests were done using a non-parametric, two-tailed paired *t*-test. To achieve this, we first computed a *t*-statistic for the actual coverage values in each ROI and each half of the visual field (upper vs. lower), and then randomly shuffled the vertical position labels for the coverage values across 10,000 iterations, and computed a *t*-statistic for each shuffling iteration. Then, we computed a two-tailed *p*-value by calculating the number of iterations on which the real *t*-statistic exceeded the shuffled statistic, and the number of iterations on which the shuffled t-statistic exceeded the real *t*-statistic, took the minimum of these values, then multiplied by 2. Finally, we performed FDR correction on the *p*-values across all ROIs using the Benjamini-Hochberg method, with an *α* value of 0.01 [96], also implemented using statsmodels.

### Regressing Out Category Selectivity

To determine whether visual feature selectivity was driven in part by category selectivity, we fit an encoding model (COCO-all model) that predicts each voxel’s response based on the COCO categories present at each location in the image (see *Methods: Semantic Features*). As described previously (*Methods: Model Fitting Procedure*), when fitting this model, we utilized the pRFs for each voxel that were already estimated based on the AlexNet encoding model, so fitting the category model for each voxel required fitting only a single set of weights across the 200 semantic category features. Once the model was fit, we used it to generate predicted voxel responses for all images in the dataset (both training and validation), and then subtracted the actual voxel activations from these predicted responses, to yield a residual for each voxel’s response to each image. These residuals represent the portion of voxel responses that cannot be modeled as a linear combination of category features. We then re-fit our Gabor encoding model with the residuals in place of raw voxel activation data, using the same approach described earlier (outlined in *Methods: Encoding Model Fitting*). Importantly, the exact same data splits were used when fitting both the COCO-all model and the Gabor model, such that the validation set data did not contribute to training of either model.

When analyzing feature selectivity results from the encoding model fit to the COCO-all residuals and comparing them to the results of the model fit to the raw data (Figure 5), we always thresholded voxels according to their R^2^ for both the raw data fit and the residuals fit, with a criterion value of 0.01 for both models. This ensured that the same set of voxels was being compared between the raw and residual fit models (see Supplementary Table 4 for the number of voxels in each ROI meeting this threshold).

### Fitting Models for Single Categories

As an additional test of whether feature selectivity was influenced by differential processing of categories, we performed Gabor model fitting using images from only one high-level category at a time (Figure 6). For this analysis, we focused on only one semantic dimension at a time, for instance, when separating images into indoor and outdoor, we ignored the animacy and real-world size labels. This was because balancing across multiple dimensions resulted in too few images to robustly fit the model. When splitting the images based on each dimension, we always split images based on labels on a per-pRF basis, meaning that the images assigned to a given category were different depending on which pRF was currently of interest. For example, when fitting the set of voxels whose best pRF was pRF *n*, we used the category labels for pRF *n* to split the images into category groupings, and performed fitting for these voxels using the data in each split separately. For a set of voxels with a different pRF estimate, the set of images that went into each split could be different. The exception to this was the indoor versus outdoor distinction, where entire images had the same label, and thus the same data split was used for all pRFs (and thus all voxels).

Given that the amount of data used to fit the encoding model can influence its overall predictive power, we always used the same number of images in the splits corresponding to a given semantic dimension and pRF. For example, if there were 5,000 animate labels for a given pRF and only 4,000 inanimate labels, we randomly selected 4,000 images from the set with the animate label. To provide an additional comparison, we also generated a subsampled set of images for each semantic dimension that was balanced with respect to category (i.e., 50 percent each label) and had the same total number of images as the smaller category (in this example, it would include 2,000 animate-labeled images and 2,000 inanimate-labeled images). Thus, for a given voxel and a given semantic dimension, the single-category and balanced category models were always fit with the same number of images, facilitating a balanced comparison between the three sets of results (i.e., indoor only, outdoor only, balanced indoor-outdoor). However, the number of images was not necessarily matched for voxels with different pRFs, or for splits over different semantic dimensions.

## Supplementary Material

**Supplementary Figure 1:**
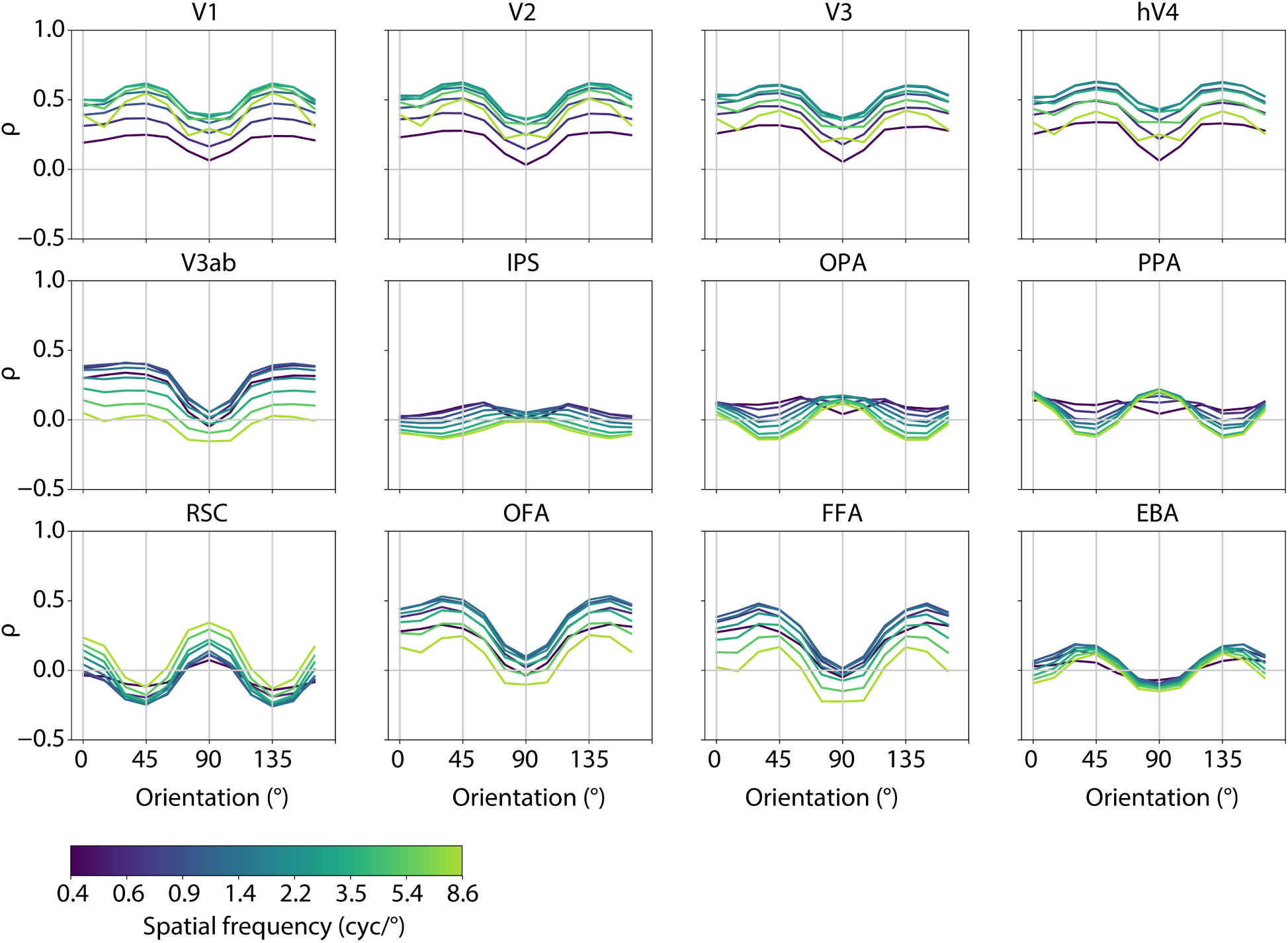
Orientation sensitivity profiles, averaged over voxels in each ROI, for the full set of 12 ROIs. Feature sensitivity (*ρ*) was estimated by computing the correlation between the predicted encoding model response for validation set images and the activation in each feature channel for the same images, see *Methods: Estimating Voxel Visual Feature Selectivity* for details. Results are averaged over all 8 participants, colored lines indicate different spatial frequency levels.

**Supplementary Figure 2:**
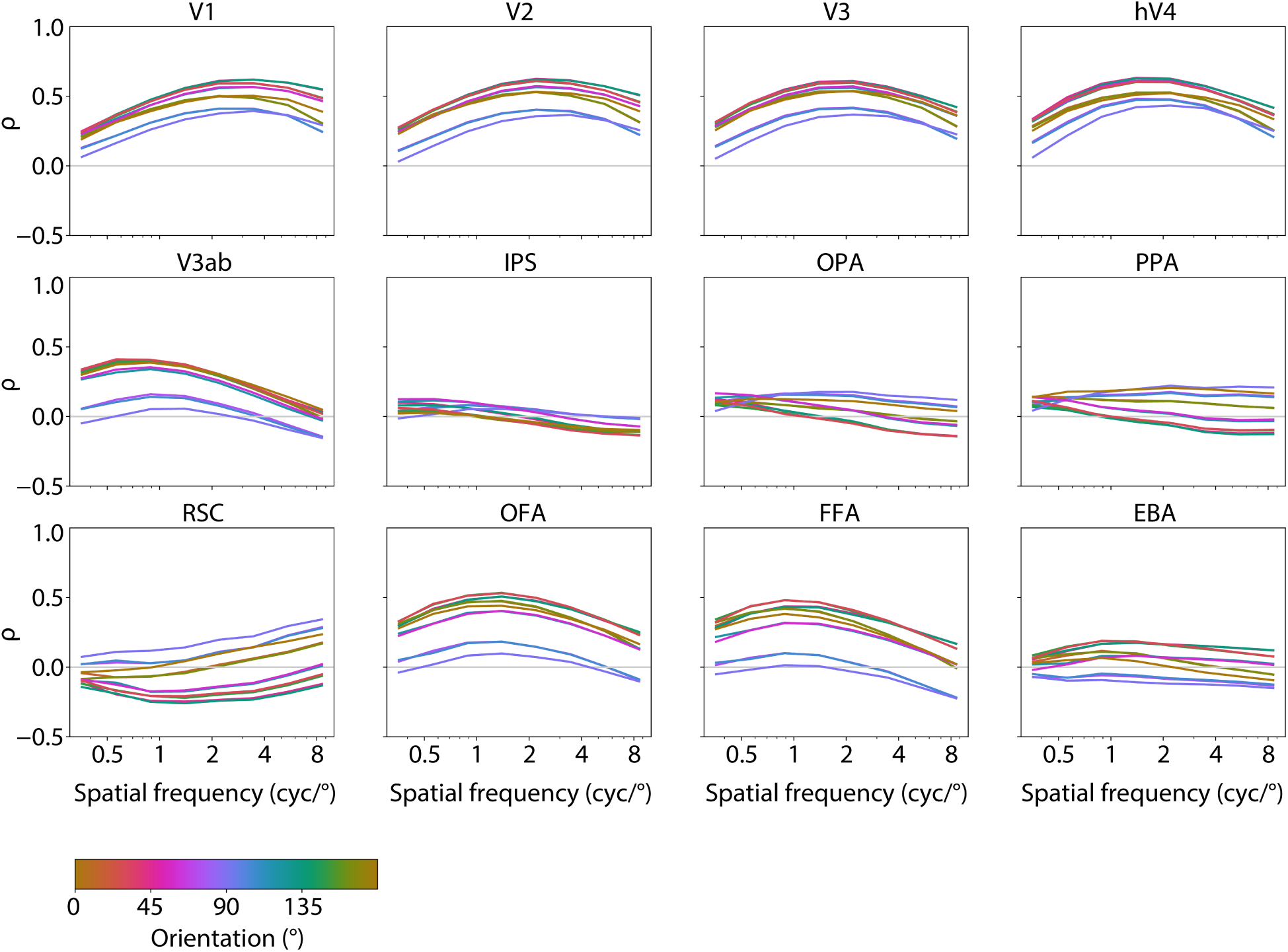
Spatial frequency sensitivity profiles, averaged over voxels in each ROI, for the full set of 12 ROIs. Feature sensitivity (*ρ*) was estimated by computing the correlation between the predicted encoding model response for validation set images and the activation in each feature channel for the same images, see *Methods: Estimating Voxel Visual Feature Selectivity* for details. Results are averaged over all 8 participants, colored lines indicate different orientations.

**Supplementary Figure 3:**
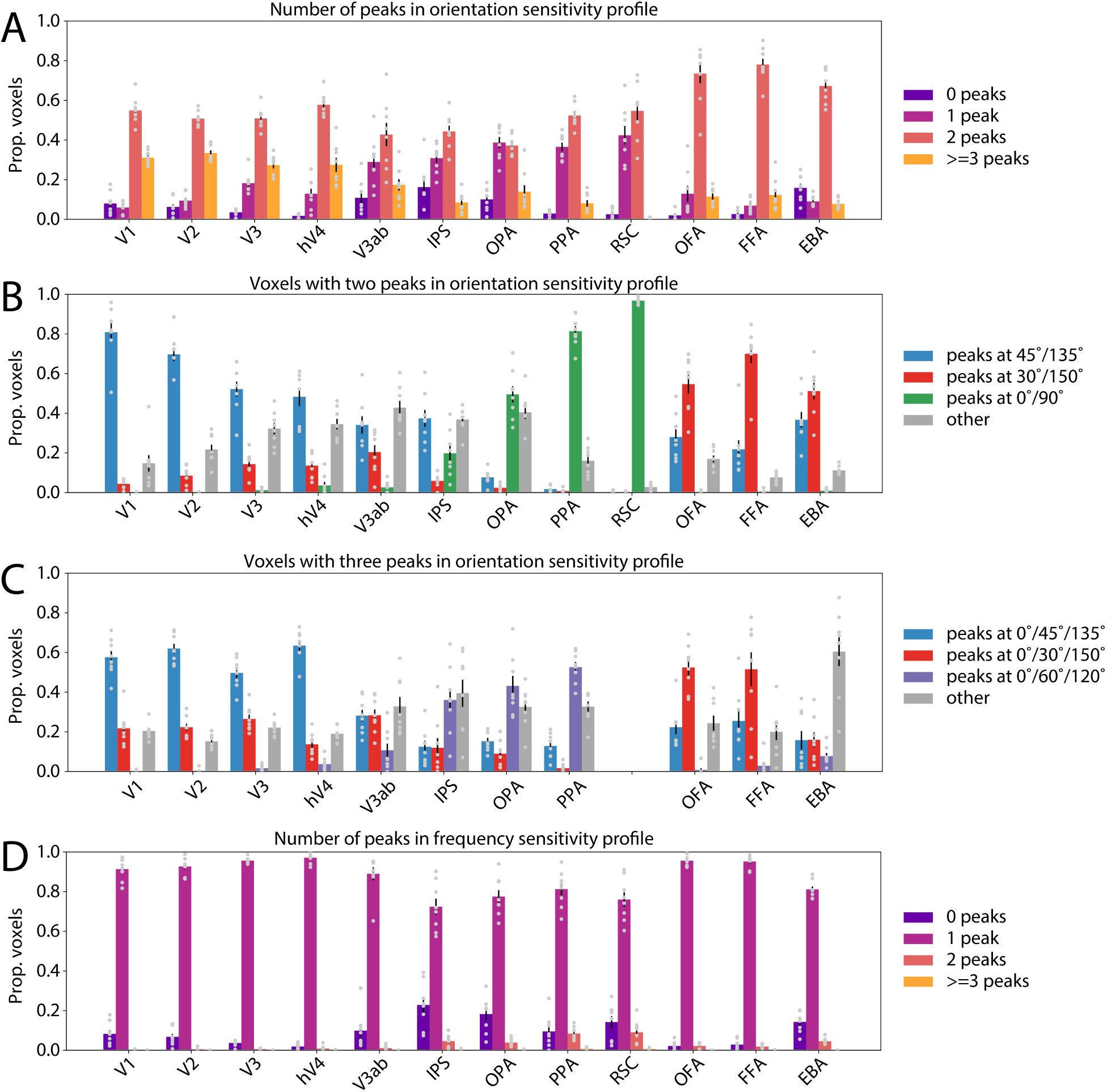
Analysis of the number of peaks in feature sensitivity profiles for each voxel, and the distribution of preferred features for multi-peaked voxels. **(A)** The proportion of voxels per ROI that had either no true peaks (zero), one peak, two peaks, or three or more peaks (see *Methods: Counting Peaks in Feature Sensitivity Profiles* for details on how peaks were counted). **(B)** The distribution of preferred orientation for voxels having exactly two peaks. Voxels were grouped based on the pair of orientations at which the two peaks occurred, without regard to the relative height of the peaks (see *Methods: Counting Peaks in Feature Sensitivity Profiles*). **(C)** The distribution of preferred orientation for voxels having exactly three peaks. RSC is excluded from this analysis because several participants did not have any three-peaked voxels in RSC. **(D)** The proportion of voxels per ROI having each number of peaks in their spatial frequency sensitivity profile. In all panels, error bars indicate mean ± 1 SEM across 8 participants, and gray dots indicate individual participants.

**Supplementary Figure 4:**
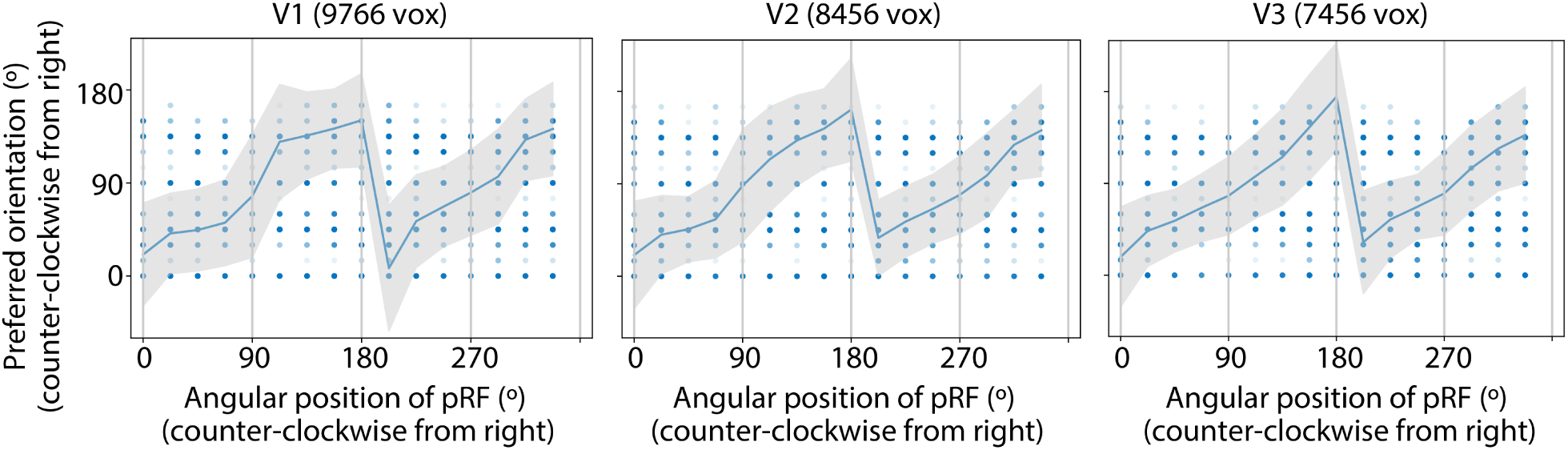
Estimates of orientation preference and angular spatial position preference are correlated across voxels in early visual ROIs, consistent with past work on the “radial bias”. Dots on each plot indicate the preferred orientation and preferred position (in polar angle) for individual voxels, where the intensity of the dot indicates the number of voxels with each preference. Blue line and shaded error bars indicate the circular mean ± 1 circular standard deviation of the preferred orientation values across all voxels sharing a given polar angle preference. Voxels in each ROI are pooled across all 8 participants.

**Supplementary Figure 5:**
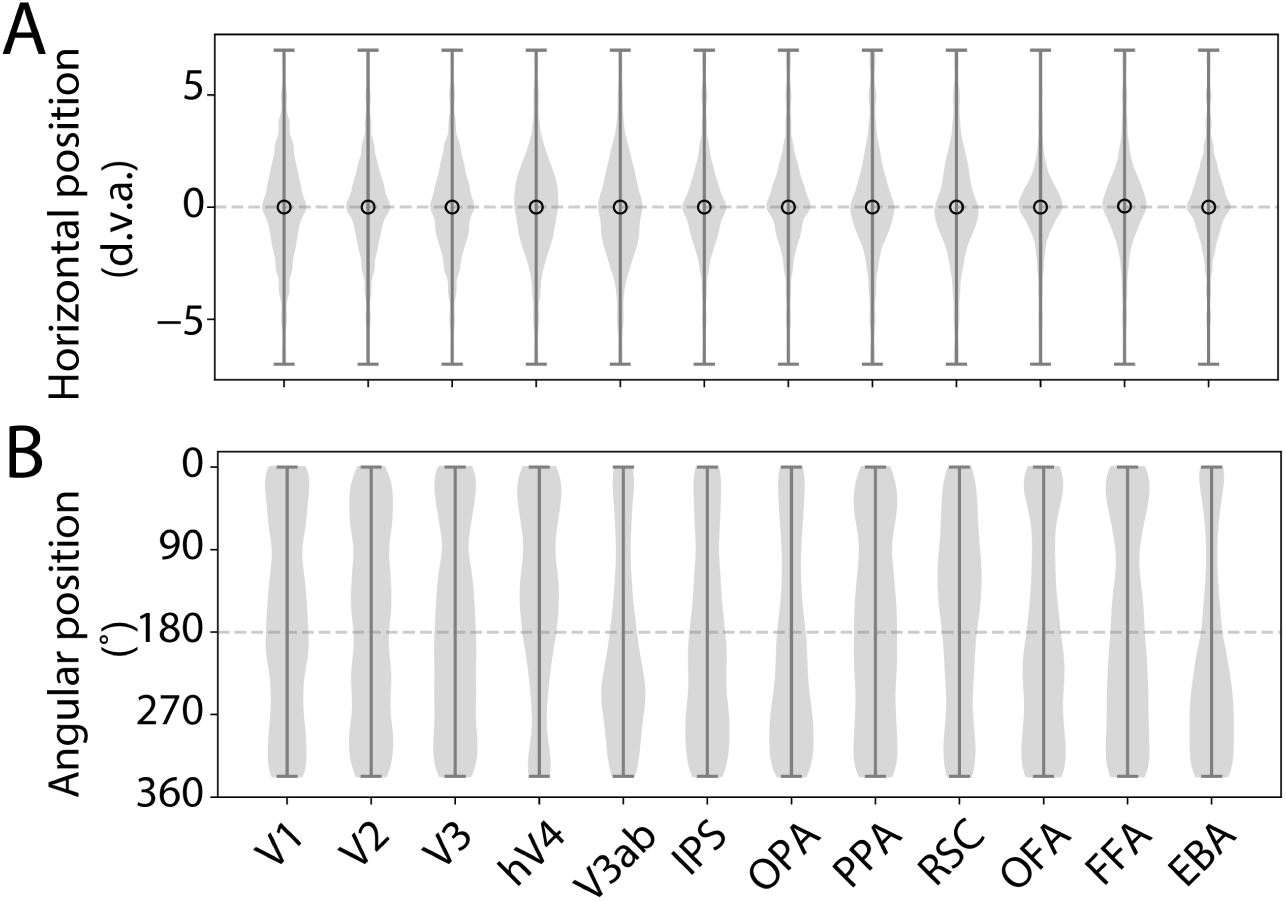
Distributions of estimated **(A)** horizontal position and **(B)** angular position preferences across voxels in each ROI. Horizontal position values are plotted from left (negative values) to right (positive values). Open circle in **(A)** indicates the median. Angular position goes in a counter-clockwise direction from the right horizontal meridian (0°), so the top of the plot corresponds to the upper visual field, and the bottom of the plot corresponds to the lower visual field. Voxels in each ROI are pooled across all 8 participants.

**Supplementary Figure 6:**
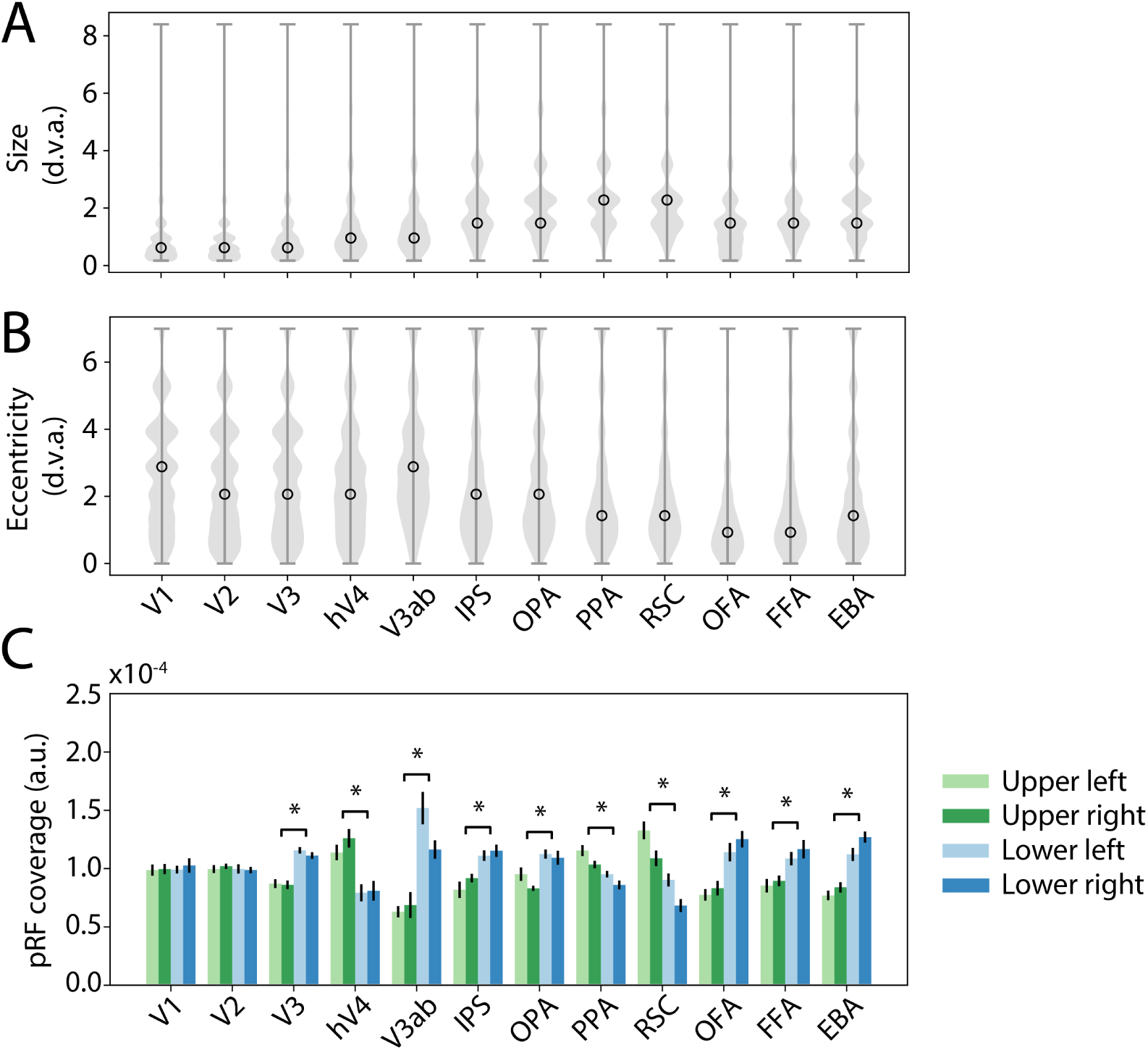
Spatial selectivity of voxels, estimated using a Gabor feature space (see *Methods: Model Fitting Procedure* for details). **(A)** Distribution of the best size (*σ*) parameter for all voxels in each ROI, combined across 8 participants. Open circle indicates the median. **(B)** Distribution of best eccentricity of pRFs across voxels. **(C)** Estimated coverage of each visual field quadrant, obtained by first aggregating pRFs across voxels in each ROI and then computing the mean of the aggregated pRF values in each visual field quadrant (see *Methods: pRF Coverage Analysis* for more details). * indicates significance of paired t-test for the upper vs. lower visual field difference, FDR corrected *α*=0.01. Errorbars indicate ± 1 SEM across participants.

**Supplementary Figure 7:**
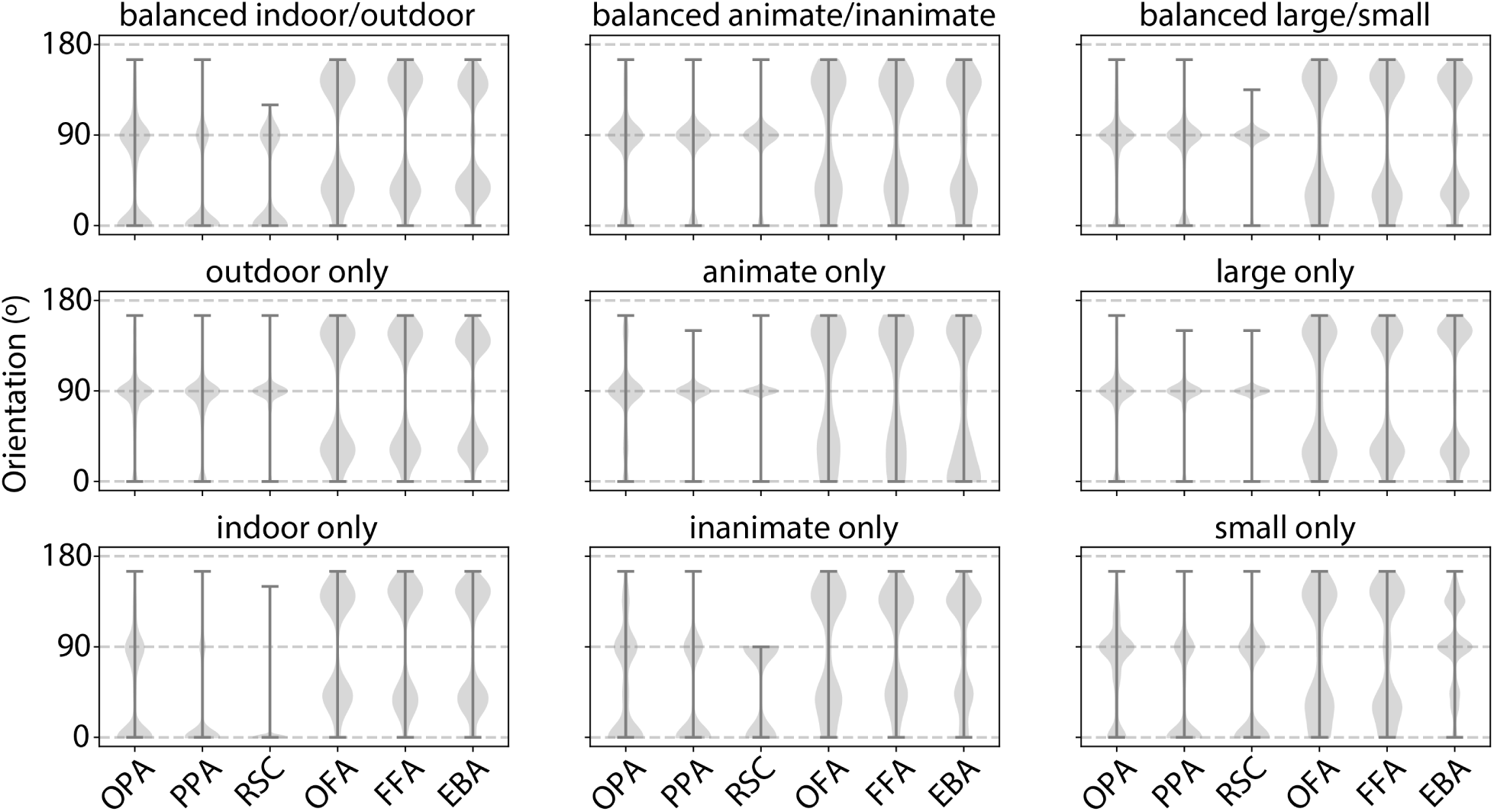
Distributions of estimated orientation preferences across voxels in each category-selective ROI, obtained by fitting encoding model on images from either one category at a time, or an image set with a balanced number of each category (see *Methods: Fitting Models for Single Categories* for details).

**Supplementary Figure 8:**
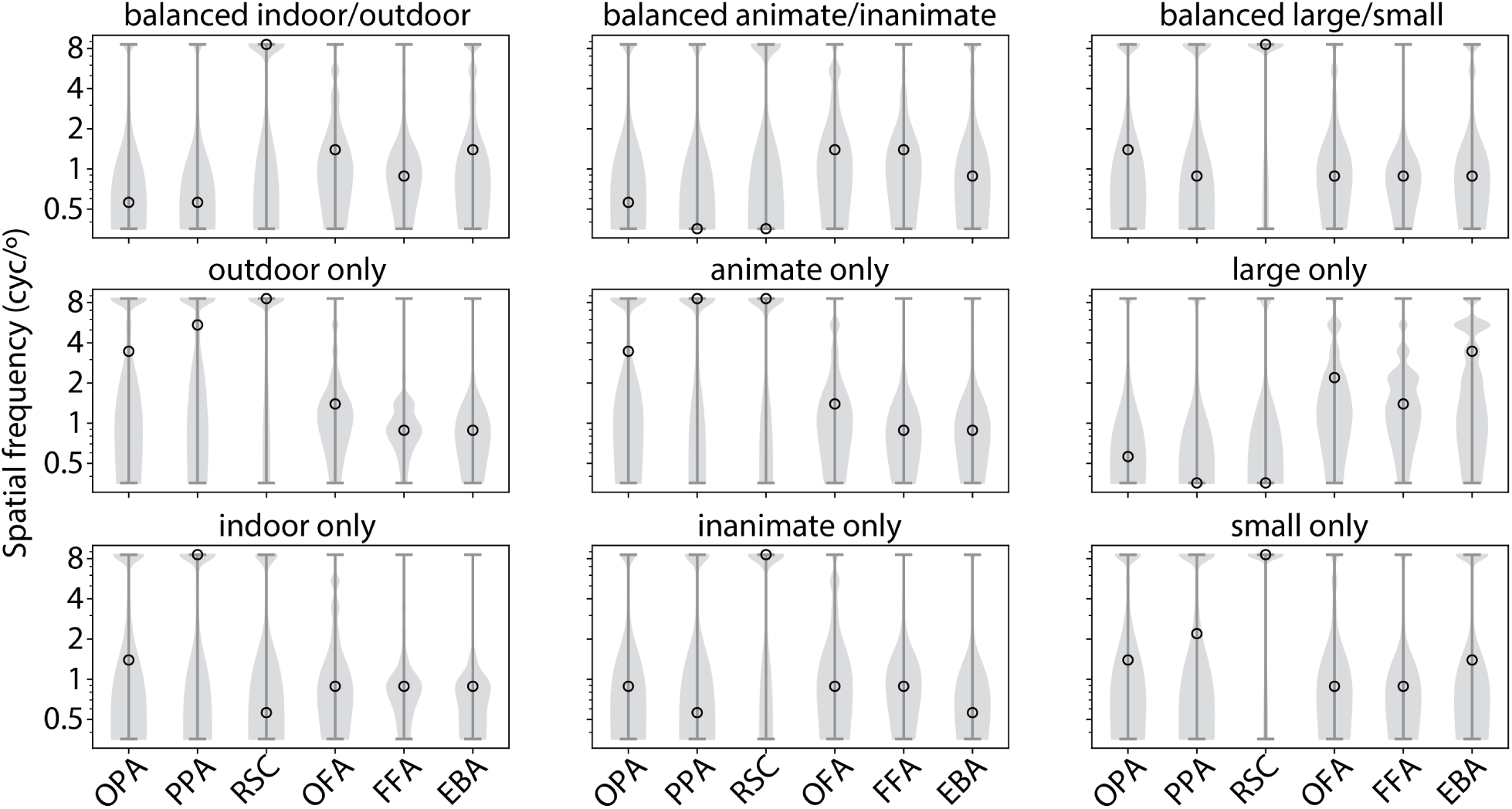
Distributions of estimated spatial frequency preferences across voxels in each category-selective ROI, obtained by fitting encoding model on images from either one category at a time, or an image set with a balanced number of each category (see *Methods: Fitting Models for Single Categories* for details).

**Supplementary Table 1:**
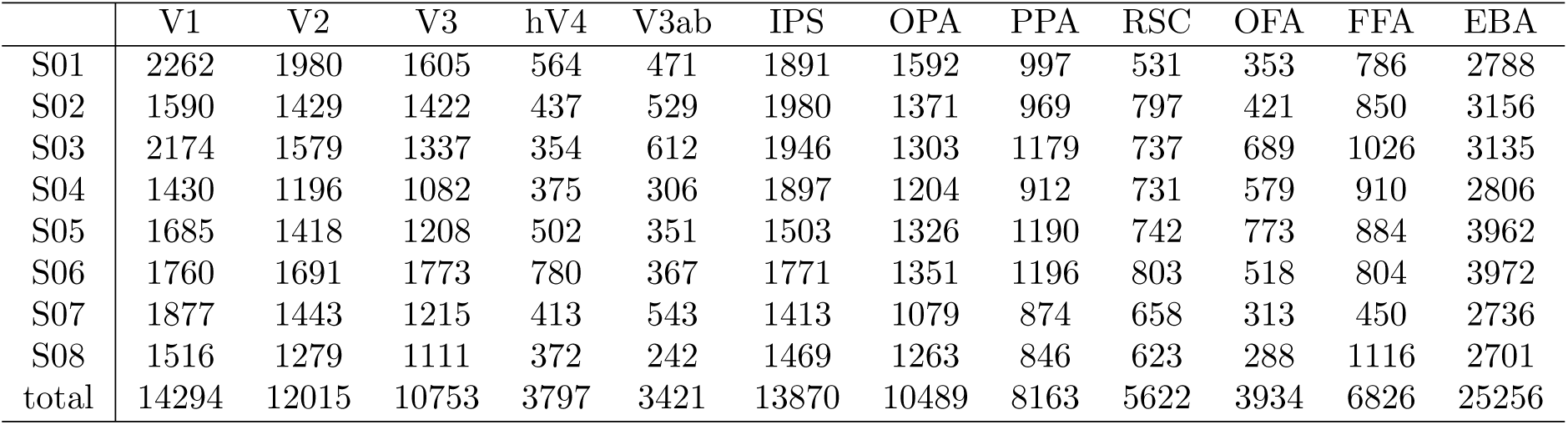
ROI sizes for all 8 participants, after thresholding by the overall noise ceiling of the voxels (see *Methods: Defining Regions of Interest (ROIs)*; noise ceiling *>* 0.01).

**Supplementary Table 2:**
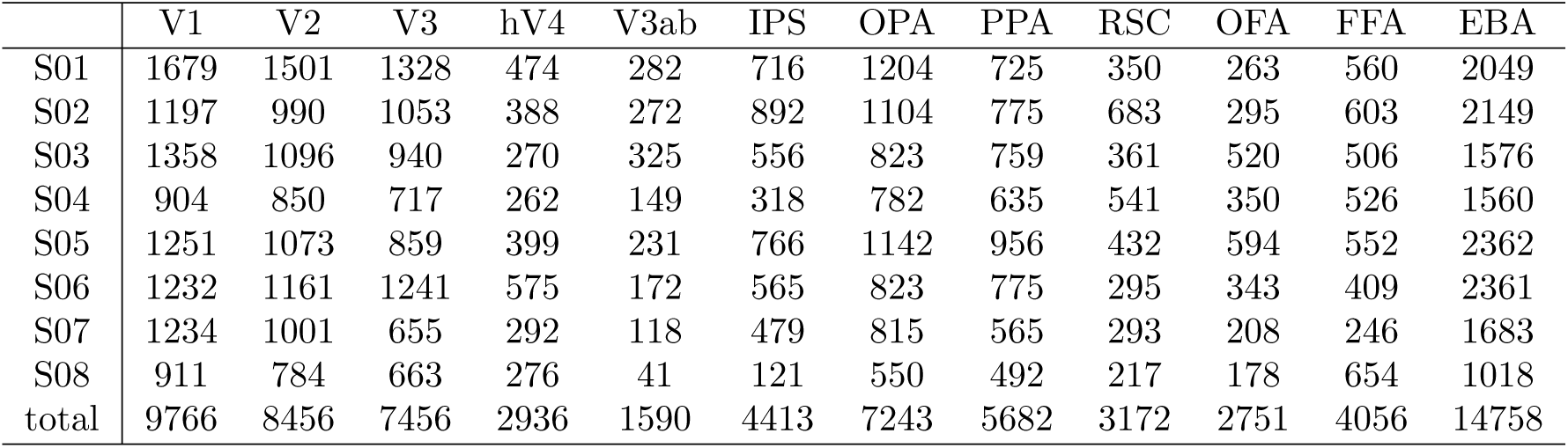
ROI sizes for all 8 participants, after thresholding by validation set R^2^ for the Gabor encoding model (R^2^ *>*0.01). This is the number of voxels used for analyses in Figure 3.

|  | V1 | V2 | V3 | hV4 | V3ab | IPS | OPA | PPA | RSC | OFA | FFA | EBA |
| --- | --- | --- | --- | --- | --- | --- | --- | --- | --- | --- | --- | --- |
| S01 | 1679 | 1501 | 1328 | 474 | 282 | 716 | 1204 | 725 | 350 | 263 | 560 | 2049 |
| S02 | 1197 | 990 | 1053 | 388 | 272 | 892 | 1104 | 775 | 683 | 295 | 603 | 2149 |
| S03 | 1358 | 1096 | 940 | 270 | 325 | 556 | 823 | 759 | 361 | 520 | 506 | 1576 |
| S04 | 904 | 850 | 717 | 262 | 149 | 318 | 782 | 635 | 541 | 350 | 526 | 1560 |
| S05 | 1251 | 1073 | 859 | 399 | 231 | 766 | 1142 | 956 | 432 | 594 | 552 | 2362 |
| S06 | 1232 | 1161 | 1241 | 575 | 172 | 565 | 823 | 775 | 295 | 343 | 409 | 2361 |
| S07 | 1234 | 1001 | 655 | 292 | 118 | 479 | 815 | 565 | 293 | 208 | 246 | 1683 |
| S08 | 911 | 784 | 663 | 276 | 41 | 121 | 550 | 492 | 217 | 178 | 654 | 1018 |
| total | 9766 | 8456 | 7456 | 2936 | 1590 | 4413 | 7243 | 5682 | 3172 | 2751 | 4056 | 14758 |

**Supplementary Table 3:** ROI sizes for all 8 participants, after thresholding by validation set R^2^ for the AlexNet encoding model (R^2^ *>*0.01). This is the number of voxels used for analyses in Figure 4.

|  | V1 | V2 | V3 | hV4 | V3ab | IPS | OPA | PPA | RSC | OFA | FFA | EBA |
| --- | --- | --- | --- | --- | --- | --- | --- | --- | --- | --- | --- | --- |
| S01 | 1709 | 1590 | 1397 | 520 | 344 | 1187 | 1428 | 851 | 403 | 304 | 648 | 2438 |
| S02 | 1261 | 1093 | 1146 | 414 | 335 | 1344 | 1236 | 841 | 713 | 344 | 686 | 2605 |
| S03 | 1526 | 1185 | 1031 | 305 | 407 | 1123 | 1056 | 898 | 391 | 586 | 721 | 2189 |
| S04 | 1022 | 952 | 855 | 327 | 201 | 845 | 1004 | 757 | 585 | 445 | 702 | 2139 |
| S05 | 1421 | 1209 | 989 | 446 | 263 | 1061 | 1247 | 1029 | 527 | 688 | 688 | 2973 |
| S06 | 1444 | 1263 | 1470 | 666 | 253 | 1011 | 1115 | 945 | 479 | 437 | 552 | 3004 |
| S07 | 1364 | 1124 | 753 | 349 | 242 | 800 | 952 | 670 | 377 | 248 | 323 | 2161 |
| S08 | 954 | 871 | 727 | 311 | 66 | 357 | 799 | 601 | 257 | 198 | 772 | 1510 |
| total | 10701 | 9287 | 8368 | 3338 | 2111 | 7728 | 8837 | 6592 | 3732 | 3250 | 5092 | 19019 |

**Supplementary Table 4:** ROI sizes for all 8 participants, after thresholding by validation set R^2^ for both the original Gabor encoding model, and R^2^ for the Gabor model fit to the COCO-all model residuals (see *Methods: Regressing Out Category Selectivity*), each at a threshold of R^2^ *>*0.01. This is the number of voxels used for analyses in Figure 5.

|  | V1 | V2 | V3 | hV4 | V3ab | IPS | OPA | PPA | RSC | OFA | FFA | EBA |
| --- | --- | --- | --- | --- | --- | --- | --- | --- | --- | --- | --- | --- |
| S01 | 1547 | 1372 | 1183 | 421 | 204 | 125 | 460 | 222 | 74 | 219 | 263 | 736 |
| S02 | 1092 | 878 | 965 | 353 | 200 | 206 | 531 | 408 | 296 | 246 | 386 | 638 |
| S03 | 1243 | 986 | 796 | 216 | 176 | 55 | 82 | 151 | 71 | 304 | 156 | 345 |
| S04 | 797 | 747 | 559 | 157 | 79 | 16 | 198 | 179 | 194 | 176 | 149 | 135 |
| S05 | 1078 | 887 | 685 | 288 | 151 | 121 | 430 | 459 | 76 | 366 | 136 | 493 |
| S06 | 1123 | 1025 | 858 | 384 | 78 | 80 | 228 | 179 | 20 | 166 | 79 | 291 |
| S07 | 1062 | 838 | 516 | 255 | 37 | 86 | 314 | 129 | 13 | 113 | 98 | 313 |
| S08 | 767 | 681 | 519 | 227 | 15 | 11 | 78 | 77 | 7 | 95 | 277 | 191 |
| total | 8709 | 7414 | 6081 | 2301 | 940 | 700 | 2321 | 1804 | 751 | 1685 | 1544 | 3142 |

**Supplementary Table 5:** Result of three-way repeated measures ANOVA on the pRF coverage values for each ROI and visual field quadrant, with factors of ROI, vertical position, and horizontal position. See *Methods: pRF Coverage Analysis* for details.

|  | F-statistic | df-num | df-denom | p |
| --- | --- | --- | --- | --- |
| roi | -2.33 | 11 | 77 | 1.0000 |
| vert | 3.29 | 1 | 7 | 0.1124 |
| horiz | 3.89 | 1 | 7 | 0.0893 |
| roi:vert | 10.68 | 11 | 77 | 0.0000 |
| roi:horiz | 0.88 | 11 | 77 | 0.5607 |
| vert:horiz | 0.07 | 1 | 7 | 0.8055 |
| roi:vert:horiz | 0.72 | 11 | 77 | 0.7191 |

